# An enormous *Paris polyphylla* genome sheds light on genome size evolution and polyphyllin biogenesis

**DOI:** 10.1101/2020.06.01.126920

**Authors:** Jing Li, Meiqi Lv, Lei Du, A Yunga, Shijie Hao, Yaolei Zhang, Xingwang Zhang, Lidong Guo, Xiaoyang Gao, Li Deng, Xuan Zhang, Chengcheng Shi, Fei Guo, Ruxin Liu, Bo Fang, Qixuan Su, Xiang Hu, Xiaoshan Su, Liang Lin, Qun Liu, Yuehu Wang, Yating Qin, Wenwei Zhang, Shengying Li, Changning Liu, Heng Li

**Author notes:** These authors contributed equally to this work (Jing Li, Meiqi Lv, Lei Du, A Yunga, Shijie Hao). Correspondence authors: Heng Li, Changning Liu, Shengying Li and Wenwei Zhang.

## Abstract

The monocot family Melanthiaceae with varying genome sizes in a range of 230-fold is an ideal model to study the genome size fluctuation in plants. Its family member *Paris* genus demonstrates an evolutionary trend of bearing huge genomes characterized by an average c-value of 49.22 pg. Here, we report a 70.18 Gb genome assembly out of the 82.55 Gb genome of *Paris polyphylla* var. yunnanensis (PPY), which represents the biggest sequenced genome to date. We annotate 69.53% repetitive sequences in this genome and 62.50% of which are long-terminal repeat (LTR) transposable elements. Further evolution analysis indicates that the giant genome likely results from the joint effect of common and species-specific expansion of different LTR superfamilies, which might contribute to the environment adaptation after speciation. Moreover, we identify the candidate pathway genes for the biogenesis of polyphyllins, the PPY-specific medicinal saponins, by complementary approaches including genome mining, comprehensive analysis of 31 next-generation RNA-seq data and 55.23 Gb single-molecule circular consensus sequencing (CCS) RNA-seq reads, and correlation of the transcriptome and phytochemical data of five different tissues at four growth stages. This study not only provides significant insights into plant genome size evolution, but also paves the way for the following polyphyllin synthetic biology.

## Introduction

*Paris polyphylla* var. yunnanensis (PPY) is a member of the genus of *Paris* in the monocot family Melanthiaceae. *Paris*, which was first described by Linnaeus in 1753, is composed of approximately 24 species, and distributed throughout Europe and East Asia, with a center of diversity in southwest China (19 species)^1^. Some *Paris* species with a thick rhizome are rich in steroidal saponins, and have long been used as medicinal herbs in traditional Chinese medicine. Among them, PPY is officially included in the Chinese Pharmacopoeia, owing to its analgesic, hemostatic, anti-tumor and anti-inflammation activities derived from the PPY-specific saponins, namely, polyphyllins^2^. Currently, there are more than 40 commercial drugs in China using the PPY rhizomes as raw materials. The wild PPY populations have been pushed to the brink of extinction due to overharvesting and habitat destruction, and therefore PPY is listed as vulnerable by the International Union for Conservation of Nature.

The genus *Paris* is subject to the Parideae tribe of Melanthiaceae which can be divided into five tribes in more than 20 genera. Notably, all the species in Parideae tribe present large-to-gigantic genome sizes according to flow cytometry analysis, and the average genome size of *Paris* species is approximately 49.22 pg (1c-value, 1 pg = 978 Mb)^3^. By contrast, species in other four tribes show much smaller genomes, varying from 0.66 pg in *Schoenocaulon texanum* to 5.50 pg in *Ypsilandra cavaleriei*^3^. The genome dimensions of the species in Melanthiaceae vary 230-fold, making it an ideal model for understanding the genome size fluctuation in plants. For the species with huge genomes in Parideae tribe, their stem age and crown age have been reported to be approximately 59.16 million years ago (MYA) and 38.21 MYA, respectively^4^. Therefore, the massive genome expansion events in Parideae have been accumulated and well-presented in over tens of millions of years, which are usually associated with transposable elements and chromosome rearrangements.

In this study, we assembled and annotated the 70.18 Gb out of the 82.55 Gb of PPY genome, which represents the biggest sequenced genome to date. We analyzed the genome sequences to investigate the possible reasons for the huge PPY genome. Furthermore, we proposed the candidate pathway genes involved in polyphyllin biosynthesis by integrative analysis of multi-omics data derived from five different tissues at four growth stages. This study sets a precedent to assemble a giant genome, providing insights into the genome size shocks during the plant evolutionary process, and lays the foundation for further researches on polyphyllin synthetic biology.

## Results and discussion

### Genome assembly and annotation

We sequenced 10.25 Tb massive parallel sequencing (MPS) reads from a 350-bp library and three mate-pair libraries, together with 1.79 Tb linked-reads from 10X Genomics Chromium technology using BGISEQ-500 sequencing platform (**Supplementary Table 1**). The genome size of PPY was estimated to be 82.55 Gb by *k*-mer analysis (**Supplementary Fig. 1** and **Supplementary Table 2**), which is far larger than the reported genome size of 53.61 Gb estimated by flow cytometry^3^. The *k*-mer distribution showed a notable signal of high repetitive sequences enriched in this genome. Because *de novo* assembly of a giant and repeat-abundant genome remains both technically challenging and cost prohibitive, development of an efficient and cost-effective assembly strategy is crucial. To overcome the bottleneck of computer memory and elevate the running time, we updated the contig module to interface the sparse *de Bruijn* graph output of the pre-graph module of SOAPdenovo2^5^ (https://github.com/BGI-Qingdao/SOAPdenovoLR). Eventually, we obtained a draft genome with the total length of 70.18 Gb (**Supplementary Table 3**), making it the largest genome assembly to date. Of note, this draft genome assembly (genome size: 82.55 Gb, scaffold N50: 21.56 Kb) is significantly improved when compared to the published giant genomes of the animal *Pleurodeles waltl*^6^ (genome size: 19.38 Gb, scaffold N50: 1.14 Kb) and the plant *Picea abies*^7^ (genome size: 19.6 Gb, scaffold N50: 4.87 Kb) using MPS data and corresponding assembly tools.

To predict the gene models of PPY genome, we collected and generated 31 MPS RNA-seq data from five different tissues (rhizome, leaf, stem, flower and fruit) at four growth stages (vegetative, flowering, fruiting and dormant stages), and 55.23 Gb single molecular PacBio CCS RNA-seq data from a sample of mixed tissues (**Supplementary Tables 4** and **5**). For the MPS RNA-seq data, we performed a series of quality control processes, such as PCA and DEG hierarchical clustering analysis (**Supplementary Figs. 2-7**). Based on these RNA-seq data, we predicted 34,257 gene models with a high complete BUSCO coverage (89.6%), revealing the good integrity and accuracy of our gene set. Briefly, 94.36% of genes were aligned against seven public databases including 83.18% in NR, 62.16% in NT, 60.15% in Swissprot, 63.79% in KEGG, 75.20% in Pfam, 64.95% in KOG and 75.36% in InterPro (**Supplementary Table 6** and **Supplementary Figs. 8-11**). According to the alignment results of NR, 32.32% and 24.01% of genes could be matched with those of *Elaeis guineensis* and *Phoenix dactylifera*, respectively (**Fig. 1a**). We reconstructed the phylogenetic tree for PPY and 11 selected species (**Fig. 1b**), and identified 3,038 expanded gene families and 3,636 contracted gene families (**Supplementary Fig. 12**). Interestingly, the enriched pathways of the expanded gene families include those involved in terpenoid biosynthesis and plant-pathogen interactions, which may play important roles in saponin biosynthesis and responses to biotic and abiotic stresses of PPY^8,9^ (**Fig. 1c** and **Supplementary Table 7**).

**Fig. 1.**
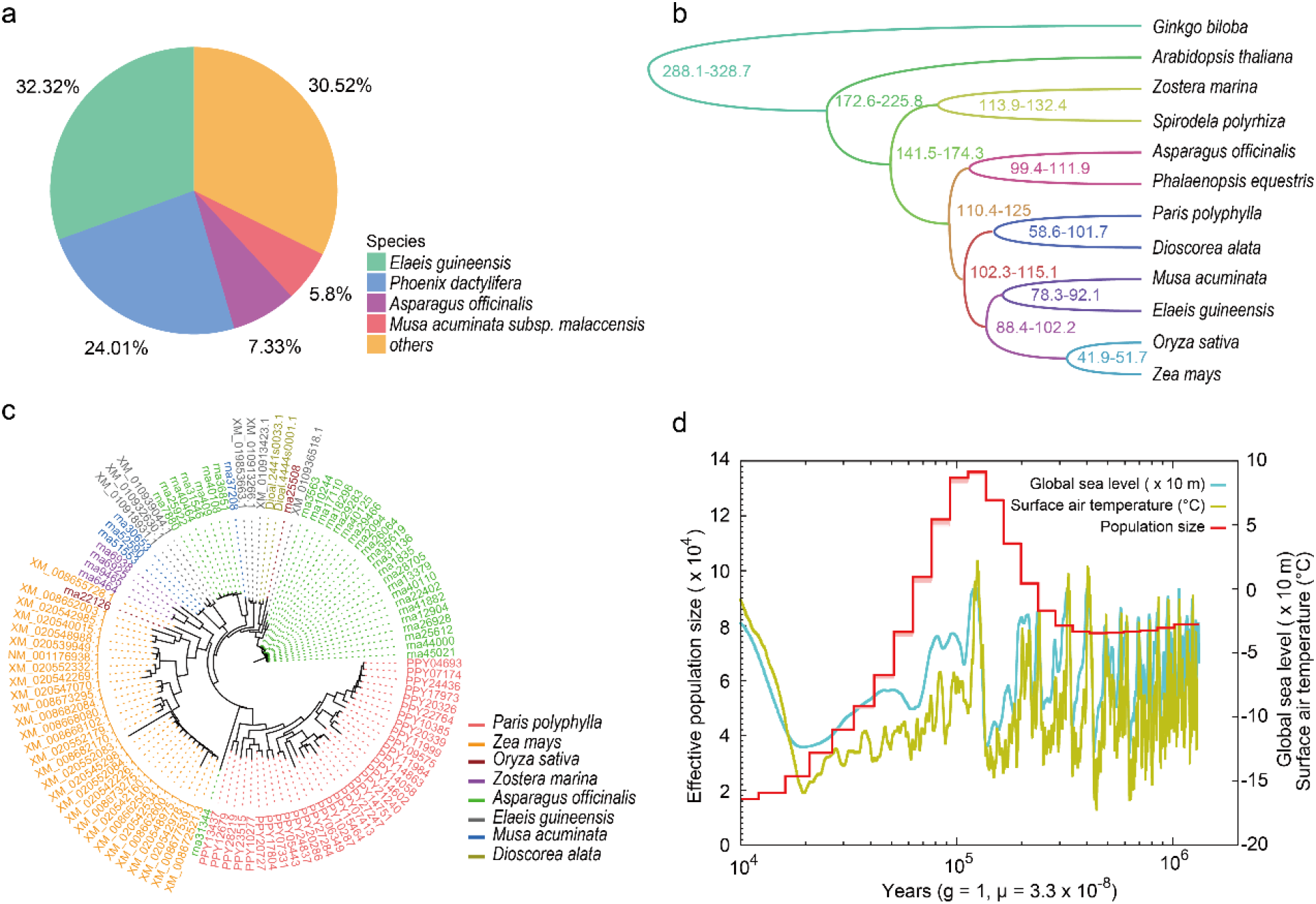
Phylogenetic analysis and genome evolution of PPY. **a**, The alignment information against NR database of the PPY gene set. **b**, The phylogenetic tree of PPY and the other eleven representative species. **c**, The gene tree of the expanded gene families related to terpenoid biosynthesis. **d**, The demographic history estimation of PPY using PSMC analysis.

### Genome size expansion and population fluctuation

The genome size diversity could have multiple causes, such as whole-genome duplication (WGD) events, and repetitive elements including transposable elements, ribosomal DNA gene copies^10^ and the abundance of centromeric repeats^11^. WGD event could also provide the foundation for the maintenance of large genome size in duplicated genomes, which is supported by both TE expansion following WGD in maize^12^, and higher TE abundance in gene-rich regions in tetraploid Capsella^13^. The species divergence time was estimated at 58.6−101.7 MYA between PPY and *Dioscorea alata* with a relatively smaller genome (**Fig. 1b**). To inquiry the candidate reasons for the enormous size of PPY genome, we detected the occurrence of WGD events in this genome by calculating the Ks values of paralogues. No obvious signal of WGD event was identified in this genome (**Supplementary Fig. 13**), suggesting the WGD event might not be a factor contributing to the PPY genome size expansion.

Next, we annotated 69.53% repetitive sequences in the assembled PPY genome, corresponding to a total of 57.86 Gb (**Supplementary Table 8**), which is lower than that of gingko (76.58%)^14^ and higher than that of maize (64.00%)^15^. Around 12 Gb of the entire genome were annotated as neither functional nor repeat sequences. These regions are most likely to be the ancient bursts of transposition followed by a long period of very low activity, resulting in a massive amount of unique sequences. We identified 50.02 Gb (62.50% of assembly) long-terminal repeat-retrotransposons (LTR-RTs), which comprises the most abundant fraction of transposable elements (TEs) (66.98% of the whole genome assembly, **Supplementary Table 9**). This is similar with those of gingko (60.65%)^14^ and maize (59.98%)^15^. Thus, the amplification of PPY genome size is mainly caused by proliferation of TEs.

The species divergence time was determined to be 48.0 MYA between PPY and its two closest phylogenetic genera *Xerophyllum*/*Heloniopsis* with relatively smaller genomes (2.5–4.75 Gb). To investigate the possible event outbursts of survival environmental changes after speciation of *Paris* and *Trillium*, we reconstructed the demographic history of PPY genome using PSMC analysis^16^. The PSMC distribution showed that PPY might have experienced a sharp decline into the rock bottom of effective population size from 100,000 to 10,000 years ago, which coincided with the increasing of sea level and surface temperature during this time (**Fig. 1d**, data from the National Climatic Data Center at http://www.ncdc.noaa.gov/). This result suggests that the environmental change could be an important factor for the effective population size of PPY.

Furthermore, we summarized the genome size, percentage of TEs and LTRs in 90 selected plant genomes that cover the major phylogenetic positions (**Fig. 2**). It was revealed that the majority of the huge genomes locates in gymnosperm, except *Paris* and *Trillium* of Melanthiaceae in angiosperms. Almost all large genomes (> 5 Gb) contain a high proportion of TEs and LTRs, which is the major reason for the large genomes of gingko, Norway spruce and others^7,14^. However, the proportion of TEs in common genomes (< 5 Gb) are unbiasedly distributed and independent of genome sizes. For example, *Cenchrus americanus* (1.58 Gb genome)^17^ and maize (2.5 Gb genome)^15^ contain a high proportion of TEs of 77.20% and 69.06%, respectively.

**Fig. 2.**
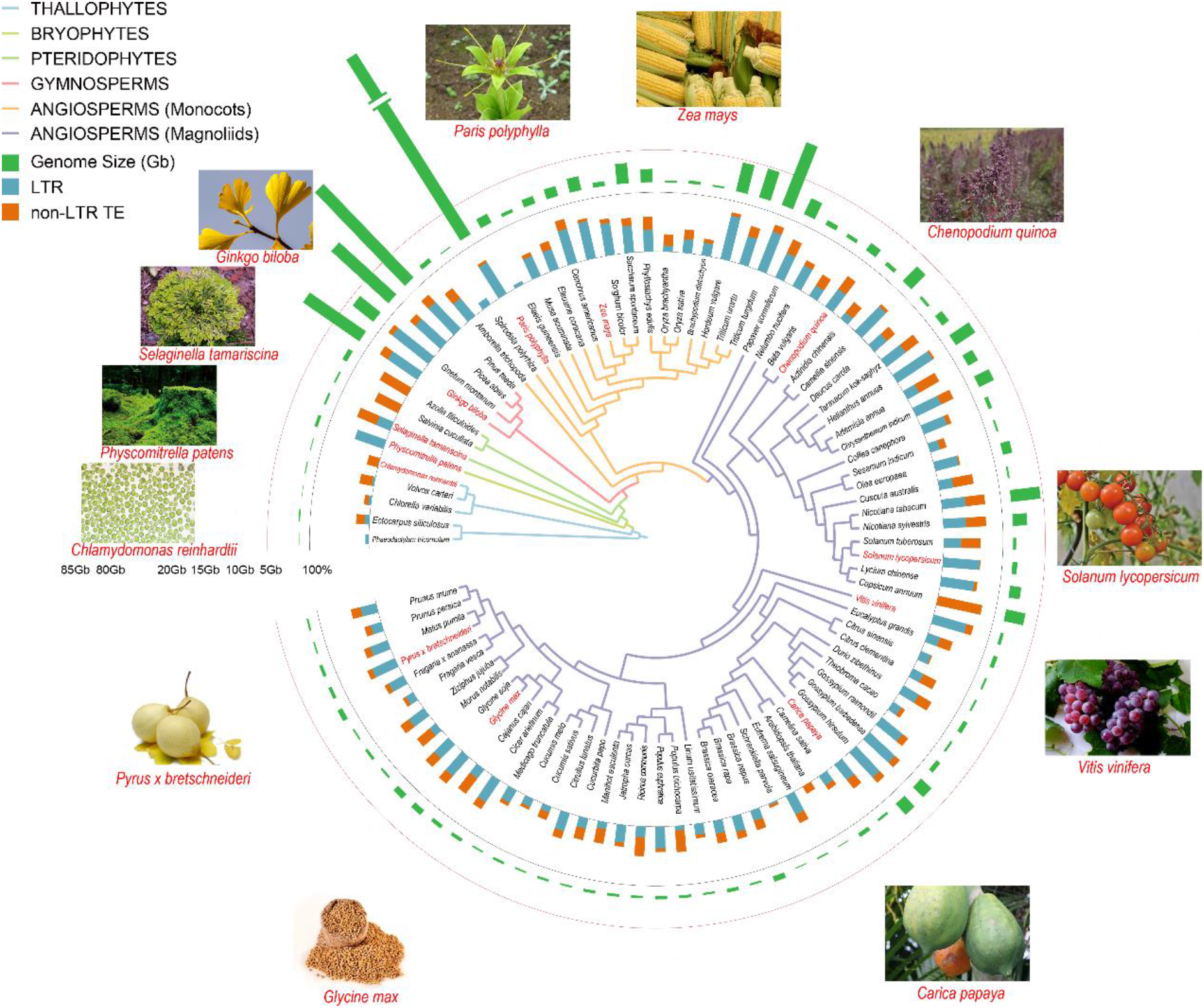
Comparison of the genome characters of representative plants. The phylogenetic tree illustrates the six main taxonomic lineages of plants using different clade colours, and the pictures of representative species in different lineages are posted. The inner circle histogram represents the percentage of LTR and non-LTR TEs in each individual genome, whereas the histogram in the outer circle denotes the genome size of each species.

### The evolution of LTR-RT in PPY genome

To trace the evolution process of LTR-RTs in PPY genome, we selected other four representative species (ginkgo, maize, rice and Arabidopsis) to perform comparative analysis. In the LTR-RTs, Ty3/Gypsy superfamily (45.69%) is more abundant than Ty1/Copia superfamily (9.99%) (**Supplementary Fig. 14**). Thus, we built the phylogenetic tree of Ty3/Gypsy and Ty1/Copia using the domains of reverse transcriptase genes. Ty3/Gypsy could be arbitrarily classified into three clades and each clade contains the Ty3/Gypsy of gingko (**Fig. 3a**), suggesting the Ty3/Gypsy of PPY genome might evolve from an ancient gymnosperm. We also found the notable expansion of three Ty3/Gypsy clades in PPY genome comparing to other four species. Calculations of the insertion times of these three Ty3/Gypsy clades indicated that two burst times (the left bulge region and the main peak region) of clade 1, clade 2 and clade 3 were around 2.2 MYA and 10.5–11.3 MYA, respectively. This suggests that all Ty3/Gypsy subtypes should have experienced two expansions (**Fig. 3b**). Moreover, we identified an earlier expansion event in clade 1 that occurred at ~18.0 MYA, revealing that most of the Ty3/Gypsy sequences burst out at a similar time.

**Fig. 3.**
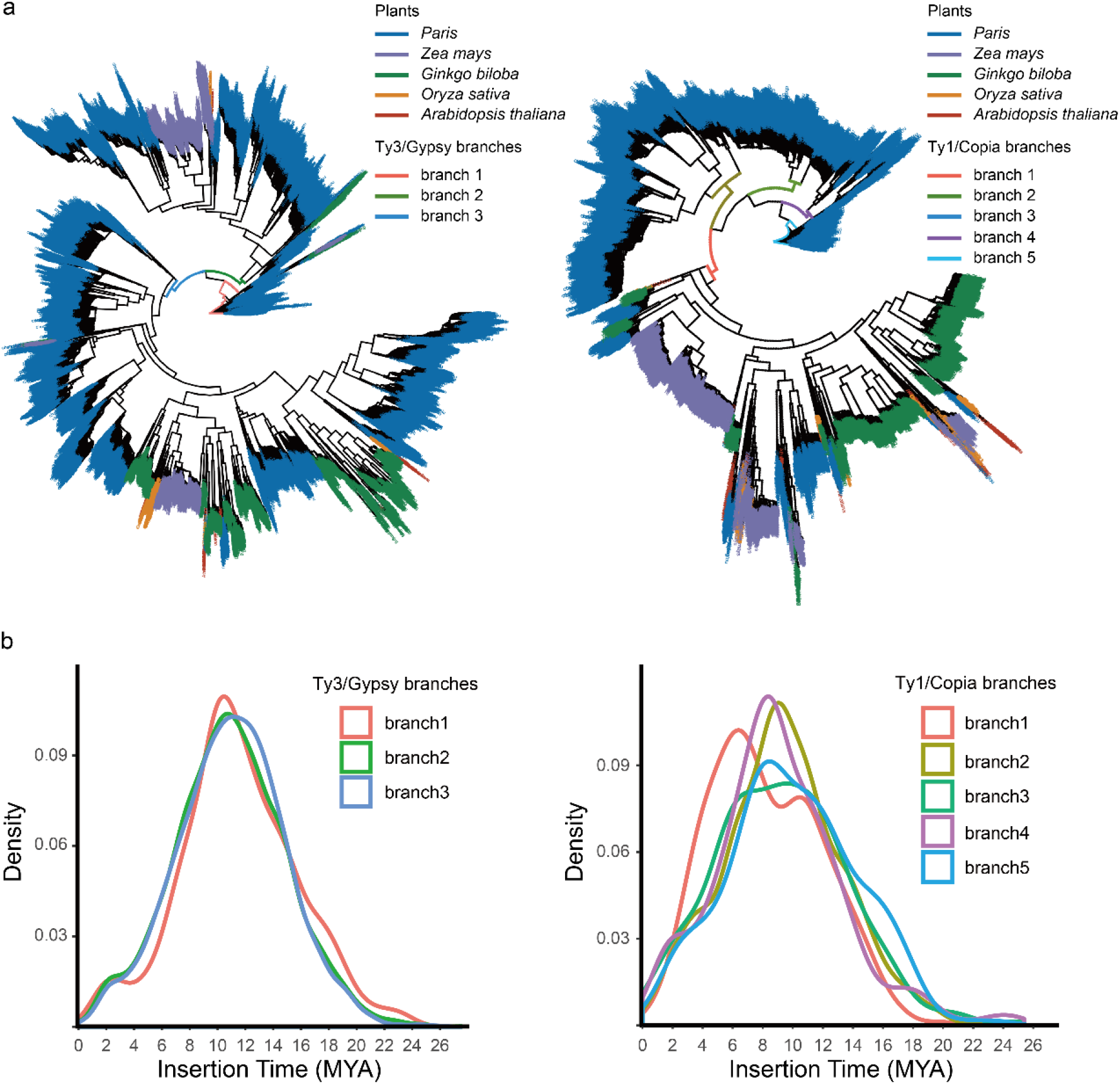
The evolution of two main subtypes of LTR. **a**, The phylogenetic trees of Ty3/Gypsy and Ty1/Copia superfamily of maize (purple), Arabidopsis (wine red), rice (yellow), ginkgo (green) and PPY (blue). **b**, The insertion times of Ty3/Gypsy and Ty1/Copia superfamily. The different coloured curves represent the insertion times of different branches in the phylogenetic tree.

For Ty1/Copia, we classified it into five clades including four species-specific amplification bursts in PPY genome and one clade shared by all the representative species at different phylogenetic positions (**Fig. 3a**). We also calculated the insertion times of these five clades. Specifically, five Ty1/Copia outburst events including the recent outburst occurred at ~2.0–3.5 MYA, the second at ~6.5 MYA, the third at ~8.4–9.1 MYA, the fourth at ~9.7–10.5 MYA, and the earliest one at ~16.0–18.0 MYA (**Fig. 3b**). Three of these events overlap with the burst time of Ty3/Gypsy, except the second and third events, suggesting PPY genome might have experienced two specific outburst events of Ty1/Copia after the common largest outburst event of Ty1/Copia and Ty3/Gypsy. For the four PPY-specific Ty1/Copia clades, the earliest outburst events in clade 4 and 5 and the largest outburst events occurred in them were detected to be ~8.4–9.1 MYA, which are earlier than the largest outburst event of the common clade (~6.5 MYA). Taken together, the giant genome size of PPY likely resulted from the common burst of three Ty3/Gypsy clades and four species-specific expanded Ty1/Copia, which might have played a critical role in the responsive capacity of PPY when facing dramatic environmental challenges in its evolutionary history^18^.

### Gene families involved in polyphyllin biosynthesis

Diosgenin and pennogenin (*i.e.*, C17α-OH-diosgenin) are the two polyphyllin aglycons with the latter being PPY-specific (**Fig. 4a**). Polyphyllin productions of 31 collected samples from five PPY tissues (rhizome, leaf, stem, flower and fruit) spanning a four-stage life cycle (vegetative, flowering, fruiting and dormant stages) were quantitatively determined by high-performance liquid chromatography (HPLC). Five saponins (*i.e.*, polyphyllins I, II, V, VII and H) were detected, among which the diosgenin-derived polyphyllins accounted for ~90% of total saponins on average, and the rest minor saponins were pennogenin-derived polyphyllins in the storage tissue rhizome (**Supplementary Table 10**). Despite the recent elucidation of diosgenin biogenesis^19^, the genes responsible for biosynthesis of pennogenin and various glycosylations to form multiple polyphyllins remain unknown. Diosgenin was previously confirmed to be produced by a number of cytochrome P450 enzymes (CYPs) including CYP90G4 and CYP94D108/CYP94D109/CYP72A616 via C16, C22 and C26 hydroxylation and the subsequent 5,6-spiroketalization of cholesterol^19^ (**Fig. 4a**). The extra C17α-hydroxy group in pennogenin suggests that an additional P450 may be required.

**Fig. 4.**
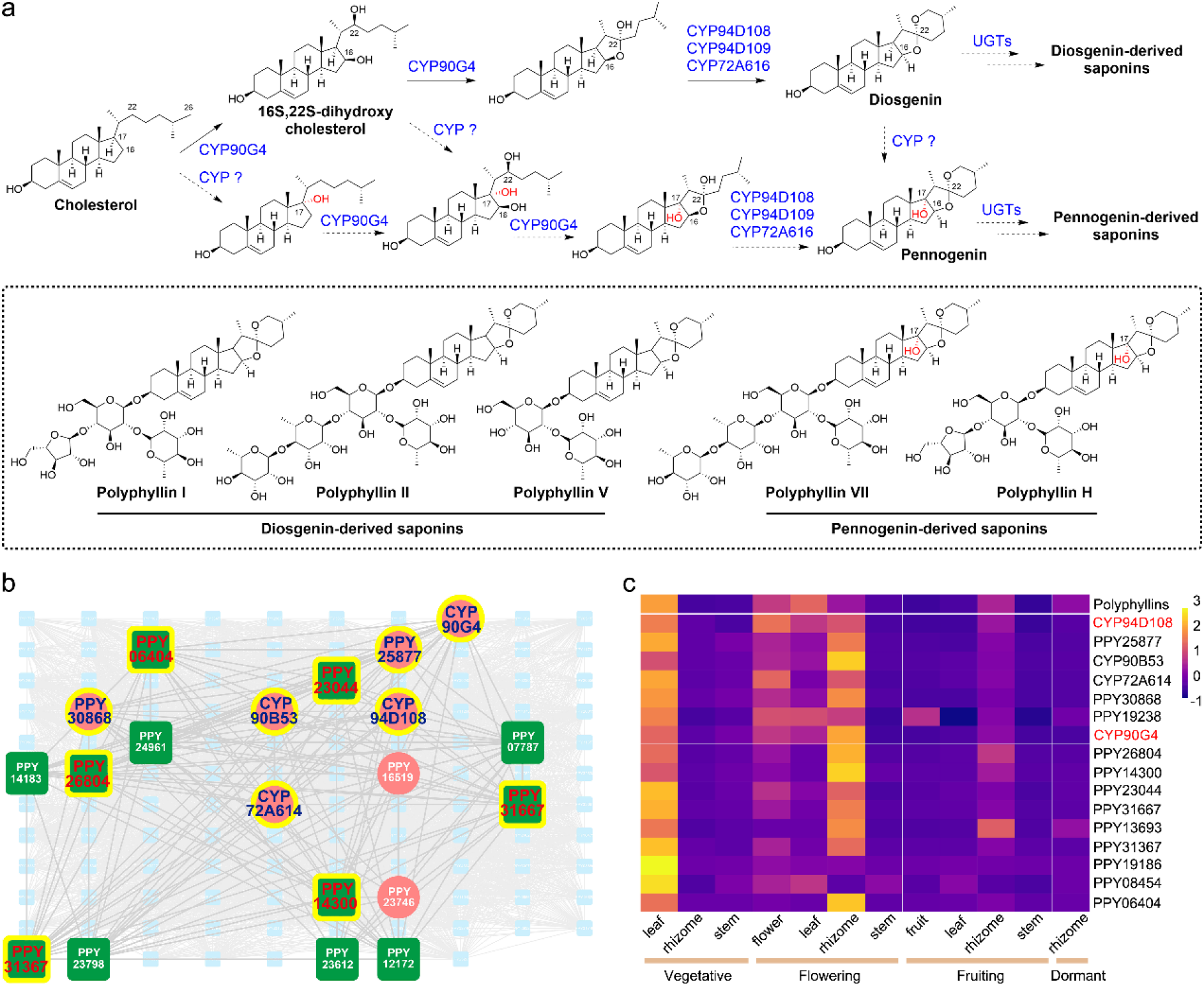
Analysis of polyphyllin biosynthesis. **a**, The proposed polyphyllin biosynthetic pathway of PPY. **b**, The co-expression network of module 35 with the P450 (red circles) and UGT (green squares) genes as well as their interactions were displayed; the P450 and UGT genes whose gene expressions are significantly correlated to the corresponding total polyphyllin concentrations are highlighted by yellow halos. **c**, The gene expression patterns of P450 (two known P450 genes related to polyphyllin biosynthesis are highlighted in red) and UGT genes that are well correlated to the total pollyphyllin production profile (*top*).

To explore the gene families involved in polyphyllin biosynthesis and their co-expression network, we performed the WGCNA analysis of 31 RNA-seq data from different PPY tissues and growth stages (**Supplementary Figs. 15** and **16**). Of the 259 identified PPY P450 genes (**Supplementary Fig. 17**), eight CYPs including CYP90G4 and CYP94D108^19^, which were proved to be involved in diosgenin biosynthesis, and other 110 genes were located in a common co-expression module (module 35) (**Fig. 4b**). These 118 genes are significantly enriched in steroid biosynthesis (ko00100, *q*-value = 4.19E-16) (**Supplementary Table 11**, **Supplementary Figs. 18** and **19**), strongly suggesting this module is closely related to biosynthesis of polyphyllins. Furthermore, transcriptome analysis of 31 RNA-seq data revealed that seven CYPs were expressed in a similar pattern that could be correlated to the total concentration of polyphyllins (**Fig. 4c**). Besides CYP90G4/CYP94D108 and another P450 enzyme CYP90B53, which is highly homologous (99% identity) to the reported CYP90B27^20^ for C-22 hydroxylation of cholesterol, there are additional three co-existing CYPs (CYP72A614, PPY25877 and PPY30868) in module 35 and the polyphyllin correlation matrix (**Fig. 4b, c**), indicative of the high possibility of these genes being involved in triterpenoid biogenesis. CYP72A614 showed a homology (62% identity) to CYP72A616^19^ for oxidative spiroketalization in diosgenin biosynthesis, while PPY25877 and PPY30868 showed homologies (80% and 82% identity, respectively) to CYP51^21^ that catalyzes the 14α-demethylation of obtusifoliol in sterol biosynthesis. Thus, we surmise that one of these three P450 enzymes might be responsible for C17 hydroxylation during pennogenin biogenesis. Biochemical confirmation of this hypothesis is currently undergoing in our laboratories.

Considering the steric hindrance of the C–H bond at C17 of diosgenin, and the fact that some isolated cholesterol-related natural products with a C17-OH retain an “open” structure without the 5,6-spiro-cyclization^22^, we hypothesized that the C17 hydroxylation is likely to occur prior to spiro-cyclization. Many plant P450s exhibit functional versatility. For example, the CYP90B subfamily has been well known to catalyze the rate-limiting sterol C22-hydroxylation in brassinosteroid (BR) biosynthetic pathway^23,24^; and it also displays low activity of sterol 16-hydroxylation^25^. Since the contents of pennogenin-derived saponins are low in the storage tissue rhizome throughout different living stages (**Supplementary Table 10**) with diosgenin-based saponins being the main components^26^, we could not exclude the possibility that pennogenin might be a “side-product” that is generated by a moonlight P450 activity during the production of diosgenin.

The late steps for polyphyllin biosynthesis require several putative UDP-dependent glycosyltransferases (UGTs). However, it is challenging to predict the specific functions of UGTs through bioinformatics analysis. For instance, UGTPg45 and PgUGT74AE2 from Panax species were reported to catalyze glycosylations at an analogous position^27^, but no homologous UGT genes could be identified in both PPY genome and transcriptomes. Of note, six UGTs were identified to co-exist in module 35 and the polyphyllin correlation matrix (**Fig. 4b, c**), which might be responsible for different polyphyllin-specific glycosylations. These genes should be good candidates for the future biochemical characterization.

## Methods

### Estimation of genome size

To estimate the genome size of PPY, the copy number of 19-mers present in approximately 30-fold clean reads from 350-bp insert size libraries was counted using kmerfreq program and the distribution of 19-mer depth of the assembled genome was plotted. The peak value in the curve represents the overall sequencing depth, and can be used to estimate genome size (G): 82.55 G = Number19-mer/Depth19-mer.

### Genome assembly

Raw data was filtered using SOAPnuke^28^ (v1.5.2) to remove the reads with PCR duplications, adaptors and a high N percentage. The original assembly version of draft genome used *SOAPdenovo2-LR* based on 42 insert size libraries data including 350 bp, 2 kb, 5 kb and 10 kb sequenced by DNBSEQ sequencing platform. The detailed parameters of *SOAPdenovo-LR* were “pregraph -s lib.cfg.1 -z -K 63 -R -d 1; contig -g -R -D 1; map -s -k 33; scaff -g out”. Next, to improve the contig continuity of the original version, we used the libraries with the insert sizes of 350 bp as the gap-filling input data of the GapCloser (v1.12, http://soap.genomics.org.cn/). To further improve the scaffold continuity, we generated ~1.4 Tb 10X Genomics sequencing data (~17.2-fold of whole genome size) with pair-end read length of 150 bp. Finally, we used ARKS^29^ to extend the scaffold and achieved the final genome assembly with default parameters.

### Gene annotation

The reads of long-read sequencing RNAseq were firstly assembled using IsoSeq (v3, https://github.com/PacificBiosciences/IsoSeq). Together with MPS RNAseq-assembled transcripts, all transcripts were deduplicated by cd-hit^30^ (v4.8.1) with strict parameters (the coverage threshold of long fragments was 0.5 while the coverage threshold of short fragments was 0.9; the sequence identity threshold was set to 0.9). Then, we predicted the open reading frames of the non-overlap transcripts with GeneMarkS-T^31^ (v5.1). Additionally, the transcripts with lengths between 500–900 nt were extracted and deduplicated with more flexible criteria (the coverage thresholds of long fragments and short fragments were both 0.5 and the sequence identity threshold was set to 0.7) using cd-hit again. The final set of transcripts was used as gene set for further analysis.

### Functional annotation of gene set

For function annotation, gene models were mapped to different functional databases, including NT (ftp://ftp.ncbi.nlm.nih.gov/blast/db), NR (ftp://ftp.ncbi.nlm.nih.gov/blast/db), GO (http://geneontology.org), KOG (http://www.ncbi.nlm.nih.gov/KOG), Pfam (http://pfam.xfam.org), KEGG (http://www.genome.jp/kegg) and SwissProt (http://ftp.ebi.ac.uk/pub/databases/swissprot). NT annotation was performed using Blastn^32^ (v2.2.23) with parameters ‘-evalue 1e-5 -max_target_seqs 5’. NR, KOG, KEGG and SwissProt annotations were performed by Diamond^33^ (v0.8.31) with parameters ‘-evalue 1e-5 -seg no -max-target-seqs 5 -more-sensitive -b 0.2’. HMMER^34^ (v 3.1b2) was used to annotate Pfam database. Finally, Blast2GO^35^ was employed to conduct GO annotation based on NR annotation results.

### Transcription factors prediction

To identify transcription factors, ORFs were detected using getorf software^36^ (EMBOSS:6.5.7.0) with the parameter ‘-minsize 150’. The predicted ORF were mapped to TF protein domains from PlntfDB^37^ database by hmmsearch with default parameters.

### Analysis of key gene families

We compared the target genes (such as CYP450) with the reference genes of *A. thaliana* using BlastP with cut-off value of 1e-5 and constructed the gene trees using MEGA software^38^. We used ClustalX^39^ (v2.1) to process multiple alignment and the protein models were predicted on the SWISS-MODEL website (https://swissmodel.expasy.org/).

### Identification of repetitive sequences and evolution

We annotated transposable elements (TEs) in the PPY genome using structure-based analyses and homology-based comparisons. Specifically, LTR_Finder^40^ (v1.0.6) and RepeatModeler (v1.0.8) were used to *de novo* identify the TEs based on features of structures. RepeatMasker (v4.0.6) and RepeatProteinMask (v4.0.6) were applied for TE identification based on RepBase^41^ at both DNA and protein levels. Finally, we double checked the overlapped TEs which were classified into the same repeat class and removed the redundant sequences. To identify Ty1/Copia and Ty3/Gypsy superfamilies in PPY genome and the genomes of the other four land plants (ginkgo, maize, rice and Arabidopsis), two-round BLAST searches were performed based on the reported Ty3/Gypsy and Ty1/Copia reverse transcriptase domains downloaded from NCBI, respectively. We used these domains to perform BLAST^32^ (v2.2.26) searches on the Ty1/Copia and Ty3/Gypsy sequences of these five land plants using parameters -p tblastn -e 1e-5 -F F -m 8, respectively. Then we extracted those sequences that satisfied the identity ≥ 0.50 and coverage ratio ≥ 0.95 as target hits. The second round of BLAST searches were performed using the target hits from round one as queries with parameter -p blastn -e 1e-5 -F F -m 8. The eventually identified amino acid sequences for all Ty1/Copia and Ty3/Gypsy elements satisfied the filter criterion of identity ≥ 0.80 and coverage ratio ≥ 0.95. To construct Ty1/Copia and Ty3/Gypsy phylogenetic trees for the five species, the resultant amino acid sequences were aligned using MAFFT^42^ (v7.245) with default parameters. Phylogenetic trees were inferred based on multiple sequence alignment using FastTree^43^.

### Estimation of effective population size

The detailed changes in the effective ancestral population size of PPY were generated using the PSMC (Pairwise sequentially Markovian coalescent) model (Version: 0.6.5-r67),^16^ which used about 30X sequencing data. We used the samtools, bcftools and vcfutils.pl (provided within the PSMC pipeline) to convert the bam file into the whole genome consensus sequence as the input file of PSMC analysis. Then PSMC was performed 50 times of bootstrap by setting parameters as follows: -N25 -t15 -r5 -b -p “4+25*2+4+6”. The number of years per generation (g) was set as one year, and the mutation rate was set as 3.3 × 10^−9^ per bp per generation.

### Calculation of the insert time of LTR-RT elements

The LTR insert time (T) was calculated based on the formula T = K / 2r^44^, where K represents the nucleotide distance between one pair of LTRs, and r represents the mutation rate. Studies reported that palm, a plant with similar evolutionary status to PPY, evolves at a rate of 2.61 × 10^−9^ substitution per synonymous site per year. Owing to mutation rates for LTR elements may be approximately 2-fold higher than synonymous substitution rates for protein-coding genes, we used a substitution rate per year of 5.22 × 10^−9^ in our calculations of LTR-RT insertion times.

### Preparation of RNA-seq tissues and *de novo* assembly

Two fruits, two flowers, eight leaves, eleven rhizomes and eight stems of PPY were collected at different developmental stages. All of these 31 samples were randomly collected from the field of greenhouse in Yunnan province, China. We carefully dealt with these samples such as cleansing, cutting into small pieces and storing in liquid nitrogen immediately after collection and then at −80 °C before library construction. We constructed the libraries according to the standard protocol of RNA-seq sequencing on DNBSEQ sequencing platform. Each cDNA library was sequenced on the DNBSEQ system using paired-end protocols. Firstly, the raw sequencing data were subjected to quality control before assembly, such as the analysis of base composition and quality filtering, removing the reads containing adapter or more than 5% low quality bases. Finally, Trinity^45^ was used for *de novo* assembly of clean reads (http://trinityrnaseq.sourceforge.net/) with default parameters. Since the raw assembled transcripts of Trinity contained redundancy, we employed TGICL^46^ and Phrap^47^ to filter the redundant data to achieve the final assembled transcripts.

### Quantitative analysis of gene expression

Clean data of RNA reads were mapped to genes by Bowtie 2^48^ (v2.2.5) with parameters ‘-q - phred33 -sensitive --dpad 0 -gbar 99999999 -mp 1,1 -np 1 -score-min L, 0, −0.1 -I 1 -X 1000 -no-mixed -no-discordant -p 1 -k 200’. Gene expression levels were calculated using RSEM software package^49^ (v1.2.12) with default parameters. Furthermore, differential expression of samples was detected by DEseq2 algorithm^50^ with the fold change ≥ 2.00 and adjusted P-value ≤ 0.05. Based on the DEG analysis results, pheatmap function in R package was used to perform hierarchical clustering analysis.

### Gene correlation network construction

The R package WGCNA^51^ (v1.69) was used to construct a gene correlation network. The offending genes and samples were filtered using *goodSamplesGenes* function. The outlier samples were filtered according to the samples tree, which was constructed by *hclust* function. Then, the network was constructed using *blockwiseModules* function. All the genes of every module were extracted, and the functional enrichment analyses were implemented respectively. The network of module 35 relating to the steroid biosynthesis, was exported to cytoscape^52^ (v3.6.1) using exportNetworkToCytoscape function. The network of module 35 was visualized by cytoscape.

### HPLC analysis of polyphyllins

The fresh samples were dried under the temperature of 40 °C, and the dried samples were smashed and sifted out with a 50 mesh sieve to obtain fine powders. Next, polyphyllins were isolated from 0.5 g of powders in a reflux system for 30 min, using 25 mL ethyl alcohol as solvent. The mixed polyphyllin standard solution containing polyphyllin I, II, V, VII and H at 0.4 mg/mL was prepared by dissolving 2 mg of each polyphyllin sample in 5 mL ethyl alcohol. The extract solutions and the standard solution were individually filtered and 10 μL of each individual sample was subjected to HPLC analysis using a Thermo HYPERSIL C-18 column with a biphasic solvent system of acetonitrile and water (30–60% acetonitrile over 40 min and 60−100% acetonitrile over 10 min; 1 mL/min; 230 nm) on an Agilent 1260 HPLC equipment.

### Correlation analysis on transcriptome and polyphyllin data

The gene expression data of all P450 and UGT genes in five different tissues at four growth stages, as well as the corresponding phytochemical data of total polyphyllin concentrations, were transformed into Z-Scores respectively. The correlation between the expression of each gene and the polyphyllin concentration was calculated by Spearman’s rank correlation coefficient, and the significantly correlated (*p*-value < 0.01) genes were extracted.

### Availability of supporting data and materials

The genome assembly and sequencing data have been deposited into CNGBdb under the project number CNP0001050. All other data are available from the corresponding authors on reasonable request.

## Acknowledgements

This work was supported by the National Key Research and Development Program (2019YFA0905100 to S.L.), the National Natural Science Foundation of China (31970609 to C.L., 31872729 S.L. and 31800041 to L.D.), Start-up Fund from Xishuangbanna Tropical Botanical Garden, ‘Top Talents Program in Science and Technology’ from Yunnan Province, the Shandong Provincial Natural Science Foundation (ZR2019ZD20 to S.L. and ZR201807060986 to L.D.), Tianjin Synthetic Biotechnology Innovation Capacity Improvement Project (TSBICIP-KJGG-001-12 to S.L.), the National Postdoctoral Innovative Talents Support Program (BX20180325 to L.D.), and China Postdoctoral Science Foundation (2019M652500 to L.D.).

## Competing interests

The authors declare that they have no competing interests.

## Author’s contributions

H.L., C.L., S.L. and W.Z. designed the project. J.L., X.G., Q.S., X.S., Y.Q., X.H., L.L., Y.W. and X.Z. prepared the samples and conducted the related experiments. L.G. and L.Deng developed the SOAPdenovo-LR tool. M.L. and Q.L. performed genome assembly. S.H. and C.S. conducted gene predictions. Y.A. and Y.Z. analyzed the repetitive sequences and carried out evolution analysis. J.L. and M.L. analyzed the polyphyllin biosynthesis related gene families. J.L., F.G., R.L., B.F. and L.Du analyzed the polyphyllin contents of the collected samples and the polyphyllin biosynthetic pathway. J.L., M.L., L.Du, Y.A., X.Z., C.L. and S.L. wrote the manuscript.

## Notes

### Competing Interest Statement

The authors have declared no competing interest.

