## Supplemental File for "An enormous *Paris polyphylla* genome sheds light on genome size evolution and polyphyllin biogenesis"

**Supplementary Table 1. Statistics of PPY genome sequencing data**

| Paired-end libraries | Insert size | Raw data (Gb) | Raw depth (X)* | Clean data (Gb) | Clean depth (X) |
|---|---|---|---|---|---|
| DNBSEQ | 350 bp | 4894.91 | 59.30 | 4767.67 | 57.75 |
| | 2 kb | 1596.89 | 19.34 | 795.22 | 9.63 |
| | 5 kb | 1638.42 | 19.85 | 476.94 | 5.78 |
| | 10 kb | 2124.70 | 25.77 | 689.03 | 8.35 |
| 10X Genomics | 350bp | 1790.81 | 21.69 | 1409.91 | 17.08 |
| Total | ---- | 12,045.73 | 145.92 | 8138.77 | 98.59 |

*The genome size (82.55 Gb) was estimated using *k*-mer analysis.

**Supplementary Table 2. The estimation of genome size using *k*-mer analysis**

| ***k*** | ***k*-mer number** | **Peak depth** | **Genome size (bp)** | **Used bases (bp)** | **X** |
|---|---|---|---|---|---|
| 19 | 2,146,347,634,136 | 26 | 82,551,832,082 | 2,617,497,114,800 | 31.71 |

**Supplementary Table 3. Summary of the genome assembly using different tools**

| | **Soap*denovo-LR*** | | **Gapcloser** | | **ARKS** | |
|---|---|---|---|---|---|---|
| | **Scaffold** | **Contig** | **Scaffold** | **Contig** | **Scaffold** | **Contig** |
| **Total number** | 15,306,712 | 78,957,901 | 15,259,092 | 57,772,429 | 15,126,348 | 57,772,429 |
| **Total length (bp)** | 70,187,646,293 | 49,921,455,841 | 68,535,337,665 | 53,296,890,466 | 68,536,665,105 | 53,296,890,466 |
| **Gap number (bp)** | 20,266,190,452 | 0 | 15,238,447,199 | 0 | 15,239,774,639 | 0 |
| **N50 length (bp)** | 20,972 | 1,814 | 20,777 | 2,400 | 21,575 | 2,400 |
| **N90 length (bp)** | 1,293 | 216 | 1,244 | 329 | 1,244 | 329 |

**Supplementary Table 4. Information of 31 RNA-seq samples**

| Sample | Time | Tissue | Sample | Time | Tissue |
|---|---|---|---|---|---|
| FF_rep1 | fruiting | fruit | PR_rep3 | flowering | rhizome |
| FF_rep2 | fruiting | fruit | PS_rep1 | flowering | stem |
| FL_rep1 | fruiting | leaf | PS_rep2 | flowering | stem |
| FL_rep2 | fruiting | leaf | PS_rep3 | flowering | stem |
| FL_rep3 | fruiting | leaf | SR_rep1 | dormant | rhizome |
| FR_rep1 | fruiting | rhizome | SR_rep2 | dormant | rhizome |
| FR_rep2 | fruiting | rhizome | SR_rep3 | dormant | rhizome |
| FS_rep1 | fruiting | stem | VL_rep2 | vegetative | leaf |
| FS_rep2 | fruiting | stem | VL_rep3 | vegetative | leaf |
| FS_rep3 | fruiting | stem | VR_rep1 | vegetative | rhizome |
| PF_rep1 | flowering | flower | VR_rep2 | vegetative | rhizome |
| PF_rep3 | flowering | flower | VR_rep3 | vegetative | rhizome |
| PL_rep1 | flowering | leaf | VS_rep1 | vegetative | stem |
| PL_rep2 | flowering | leaf | VS_rep2 | vegetative | stem |
| PL_rep3 | flowering | leaf | VS_rep3 | vegetative | stem |
| PR_rep1 | flowering | rhizome | | | |

**Supplementary Table 5. Statistics of 31 RNA-seq data from different tissues**

| Sample | Total data* (Gb) | Read length (bp) | Number of reads | Q30 of reads1 (%) | Q30 of reads2 (%) |
|---|---|---|---|---|---|
| FF_rep1 | 6.19 | 150 | 41,298,998 | 94.84 | 88.13 |
| FF_rep2 | 6.90 | 150 | 46,032,854 | 94.62 | 86.56 |
| FL_rep1 | 9.93 | 150 | 66,193,224 | 94.67 | 87.01 |
| FL_rep2 | 6.24 | 150 | 41,586,656 | 94.65 | 88.06 |
| FL_rep3 | 6.75 | 150 | 45,005,944 | 94.81 | 87.48 |
| FR_rep1 | 6.28 | 150 | 41,897,230 | 94.88 | 87.73 |
| FR_rep2 | 6.63 | 150 | 44,229,690 | 94.92 | 88.28 |
| FS_rep1 | 7.17 | 150 | 47,815,670 | 94.87 | 88.28 |
| FS_rep2 | 6.57 | 150 | 43,831,294 | 94.71 | 88.00 |
| FS_rep3 | 5.99 | 150 | 39,947,070 | 93.88 | 85.63 |
| PF_rep1 | 6.96 | 150 | 46,425,952 | 94.75 | 89.51 |
| PF_rep3 | 7.51 | 150 | 50,090,044 | 94.79 | 89.50 |
| PL_rep1 | 7.57 | 150 | 50,496,270 | 94.74 | 88.96 |
| PL_rep2 | 6.91 | 150 | 46,033,580 | 94.67 | 88.47 |
| PL_rep3 | 7.52 | 150 | 50,137,262 | 94.70 | 88.89 |
| PR_rep1 | 6.90 | 150 | 45,985,930 | 94.82 | 90.08 |
| PR_rep3 | 7.19 | 150 | 47,943,132 | 94.64 | 89.78 |
| PS_rep1 | 6.93 | 150 | 46,171,660 | 94.79 | 89.91 |
| PS_rep2 | 8.09 | 150 | 53,957,884 | 94.77 | 89.02 |
| PS_rep3 | 6.83 | 150 | 45,555,126 | 94.70 | 88.75 |
| SR_rep1 | 6.52 | 150 | 43,488,178 | 94.98 | 89.12 |
| SR_rep2 | 7.00 | 150 | 46,636,278 | 94.95 | 89.00 |
| SR_rep3 | 7.24 | 150 | 48,234,288 | 95.13 | 89.77 |
| VL_rep2 | 8.24 | 150 | 54,909,972 | 96.05 | 86.08 |
| VL_rep3 | 7.24 | 150 | 48,290,528 | 96.03 | 86.60 |
| VR_rep1 | 7.22 | 150 | 48,115,186 | 95.38 | 82.93 |
| VR_rep2 | 6.91 | 150 | 46,048,814 | 95.53 | 84.68 |
| VR_rep3 | 7.71 | 150 | 51,424,852 | 95.98 | 87.14 |
| VS_rep1 | 7.24 | 150 | 48,242,640 | 95.90 | 86.89 |
| VS_rep2 | 7.83 | 150 | 52,182,552 | 95.95 | 87.11 |
| VS_rep3 | 7.10 | 150 | 47,303,662 | 96.00 | 86.79 |

* Total size of sequenced raw data. All other column descriptions were obtained from clean data.

**Supplementary Table 6. Statistics for functional annotation of geneset**

| Values | Nr | Nt | Swissprot | KEGG | KOG | Pfam | Interpro | Intersection | Overall |
|---|---|---|---|---|---|---|---|---|---|
| Number | 28,496 | 21,295 | 20,605 | 21,853 | 22,249 | 25,762 | 25,816 | 3,845 | 32,324 |
| Percentage | 83.18% | 62.16% | 60.15% | 63.79% | 64.95% | 75.20% | 75.36% | 11.22% | 94.36% |

**Supplementary Table 7. The enriched KEGG pathways of expanded gene families**

| #Pathway | Expanded genes | All genes | P-value | Q-value | Pathway ID |
|---|---|---|---|---|---|

| | | | | | |
|---|---|---|---|---|---|
| Plant-pathogen interaction | 386 | 1131 | 2.86E-184 | 2.46E-182 | ko04626 |
| Cyanoamino acid metabolism | 101 | 261 | 1.82E-52 | 7.81E-51 | ko00460 |
| RNA transport | 256 | 1527 | 3.91E-48 | 1.12E-46 | ko03013 |
| Starch and sucrose metabolism | 137 | 591 | 5.05E-41 | 1.09E-39 | ko00500 |
| Tryptophan metabolism | 76 | 236 | 2.48E-33 | 4.27E-32 | ko00380 |
| MAPK signaling pathway - plant | 132 | 684 | 8.25E-31 | 1.18E-29 | ko04016 |
| Phenylalanine metabolism | 59 | 167 | 1.34E-28 | 1.65E-27 | ko00360 |
| Arginine and proline metabolism | 62 | 187 | 3.24E-28 | 3.49E-27 | ko00330 |
| RNA polymerase | 165 | 1094 | 1.97E-25 | 1.88E-24 | ko03020 |
| Phenylpropanoid biosynthesis | 123 | 695 | 3.36E-25 | 2.89E-24 | ko00940 |
| Purine metabolism | 177 | 1318 | 1.71E-21 | 1.34E-20 | ko00230 |
| Pyrimidine metabolism | 169 | 1252 | 8.97E-21 | 6.43E-20 | ko00240 |
| RNA degradation | 103 | 668 | 7.29E-17 | 4.82E-16 | ko03018 |
| Amino sugar and nucleotide sugar metabolism | 94 | 591 | 2.31E-16 | 1.42E-15 | ko00520 |
| Other glycan degradation | 53 | 232 | 2.82E-16 | 1.62E-15 | ko00511 |
| Aminoacyl-tRNA biosynthesis | 49 | 228 | 4.65E-14 | 2.50E-13 | ko00970 |
| Spliceosome | 104 | 819 | 1.62E-11 | 8.22E-11 | ko03040 |
| Metabolic pathways | 527 | 6415 | 1.81E-11 | 8.67E-11 | ko01100 |
| Sesquiterpenoid and triterpenoid biosynthesis | 29 | 123 | 6.79E-10 | 3.07E-09 | ko00909 |
| Glutathione metabolism | 33 | 181 | 4.61E-08 | 1.98E-07 | ko00480 |
| Diterpenoid biosynthesis | 36 | 209 | 5.26E-08 | 2.11E-07 | ko00904 |
| Alanine, aspartate and glutamate metabolism | 30 | 156 | 5.41E-08 | 2.11E-07 | ko00250 |
| Pentose and glucuronate interconversions | 61 | 474 | 1.64E-07 | 6.15E-07 | ko00040 |
| Galactose metabolism | 55 | 415 | 2.54E-07 | 9.11E-07 | ko00052 |
| Ascorbate and aldarate metabolism | 44 | 306 | 4.22E-07 | 1.45E-06 | ko00053 |
| Protein processing in endoplasmic reticulum | 138 | 1435 | 9.48E-07 | 3.14E-06 | ko04141 |
| Sphingolipid metabolism | 32 | 201 | 1.71E-06 | 5.45E-06 | ko00600 |
| Isoquinoline alkaloid biosynthesis | 17 | 89 | 4.36E-05 | 1.34E-04 | ko00950 |
| Selenocompound metabolism | 14 | 78 | 0.00039458 | 1.17E-03 | ko00450 |
| Pyruvate metabolism | 31 | 281 | 0.00227293 | 6.52E-03 | ko00620 |
| One carbon pool by folate | 8 | 44 | 0.00623706 | 1.73E-02 | ko00670 |
| Folate biosynthesis | 12 | 83 | 0.00645847 | 1.74E-02 | ko00790 |

**Supplementary Table 8. The summary of the repetitive sequences using different approaches**

| Type | #Repeat sequences (bp) | % of genome |
|---|---|---|
| Trf | 3,053,854,723 | 3.67 |
| Repeatmasker | 6,403,297,891 | 7.69 |
| Proteinmask | 9,839,396,372 | 11.82 |
| *De novo* | 56,619,598,867 | 68.03 |
| Total | 57,866,104,418 | 69.53 |

**Supplementary Table 9. Summary of the different repetitive sequence types**

| | **Repbase TEs** | | **TEs** | | **De novo** | | **Combined TEs** | |
|---|---|---|---|---|---|---|---|---|
| | Length (bp) | % in genome | Length (bp) | % in genome | Length (bp) | % in genome | Length (bp) | % in genome |
| DNA | 531,888,496 | 0.64 | 404,432,901 | 0.49 | 3,259,511,564 | 3.92 | 3,527,630,898 | 4.24 |
| LINE | 134,837,627 | 0.16 | 188,998,388 | 0.23 | 1,003,411,788 | 1.21 | 1,175,068,929 | 1.41 |
| SINE | 4,013,931 | 0.00 | 0 | 0.00 | 55,895,675 | 0.07 | 59,348,390 | 0.07 |
| LTR | 5,772,137,553 | 6.94 | 9,247,929,145 | 11.11 | 51,321,030,866 | 61.67 | 52,016,651,054 | 62.50 |
| Other | 33,931 | 0.00 | 1,563 | 0.00 | 38,889 | 0.00 | 74,383 | 0.00 |
| Unknown | 0 | 0.00 | 1,404 | 0.00 | 811,718,362 | 0.98 | 811,719,766 | 0.98 |
| Total | 6,403,297,891 | 7.69 | 9,839,396,372 | 11.82 | 54,937,887,931 | 66.01 | 55,742,660,920 | 66.98 |

**Supplementary Table 10. Pennogenin concentration in different tissues**

| **No.** | **Stage** | **Tissue** | **Diosgenin conc.** | **Pennogenin conc.** | **Diosgenin percentage** |
|---|---|---|---|---|---|
| QJPPY7R01V | Vegetative | rhizome | 0.1219 | 0.0000 | 100.00% |
| QJPPY7R02V | | rhizome | 0.1668 | 0.1325 | 55.72% |
| QJPPY7R03V | | rhizome | 0.1006 | 0.1108 | 47.60% |
| QJPPY7S01V | | stem | 0.2599 | 0.0000 | 100.00% |
| QJPPY7S02V | | stem | 0.1890 | 0.0446 | 80.92% |
| QJPPY7S03V | | stem | 0.1499 | 0.0490 | 75.37% |
| QJPPY7L01V | | leaf | 3.9549 | 0.2193 | 94.75% |
| QJPPY7L02V | | leaf | 0.9450 | 0.3699 | 71.87% |
| QJPPY7L03V | | leaf | 0.9379 | 0.0000 | 100.00% |
| QJPPY7R01P | Flowering | rhizome | 0.5019 | 0.1794 | 73.67% |
| QJPPY7R02P | | rhizome | 1.2821 | 0.0868 | 93.66% |
| QJPPY7R03P | | rhizome | 0.3460 | 0.0609 | 85.02% |
| QJPPY7S01P | | stem | 0.0000 | 0.0696 | 0.00% |
| QJPPY7S02P | | stem | 0.4094 | 0.1847 | 68.92% |
| QJPPY7S03P | | stem | 0.1826 | 0.0000 | 100.00% |
| QJPPY7L01P | | leaf | 1.7093 | 1.2425 | 57.91% |
| QJPPY7L02P | | leaf | 0.0000 | 0.3300 | 0.00% |
| QJPPY7L03P | | leaf | 0.0000 | 0.0000 | |
| QJPPY7F01P | | flower | 1.8666 | 1.1596 | 61.68% |
| QJPPY7F02P | | flower | 0.0000 | 0.3791 | 0.00% |
| QJPPY7F03P | | flower | 0.3094 | 0.0000 | 100.00% |
| QJPPYR0908A | Fruiting | rhizome | 1.0465 | 0.0892 | 92.15% |
| QJPPYR0908B | | rhizome | 0.7953 | 0.1218 | 86.72% |
| QJPPYR0908C | | rhizome | 0.8430 | 0.0784 | 91.49% |
| QJPPYS0908A | | stem | 0.0400 | 0.0000 | 100.00% |
| QJPPYS0908B | | stem | 0.0983 | 0.0279 | 77.87% |
| QJPPYS0908C | | stem | 0.0488 | 0.0543 | 47.34% |
| QJPPYL0908A | | leaf | 0.1144 | 0.0000 | 100.00% |
| QJPPYL0908B | | leaf | 0.3606 | 0.0000 | 100.00% |
| QJPPYL0908C | | leaf | 0.0000 | 0.0000 | |
| QJPPYF0908A | | fruit | 0.0507 | 0.1412 | 26.40% |
| QJPPYF0908B | | fruit | 0.0770 | 0.0756 | 50.45% |
| QJPPYF0908C | | fruit | 0.0000 | 0.0610 | 0.00% |
| QJPPYR1111A | Dormant | rhizome | 0.4857 | 0.0641 | 88.34% |
| QJPPYR1111B | | rhizome | 0.9208 | 0.0518 | 94.68% |
| QJPPYR1111C | | rhizome | 0.6978 | 0.0441 | 94.06% |

**Supplementary Table 11. The KEGG enrichment results of the genes in module 35**

| #Pathway | Genes in KO | All genes | P-value | Q-value | Pathway ID |
|---|---|---|---|---|---|
| Biosynthesis of secondary metabolites | 51 | 3081 | 1.57E-21 | 1.00E-19 | ko01110 |
| Steroid biosynthesis | 12 | 63 | 1.31E-17 | 4.19E-16 | ko00100 |
| Metabolic pathways | 53 | 6415 | 1.54E-09 | 3.28E-08 | ko01100 |
| Carotenoid biosynthesis | 6 | 107 | 3.82E-06 | 6.11E-05 | ko00906 |
| Terpenoid backbone biosynthesis | 5 | 102 | 4.87E-05 | 6.23E-04 | ko00900 |
| Synthesis and degradation of ketone bodies | 2 | 10 | 0.0006 | 6.98E-03 | ko00072 |
| Brassinosteroid biosynthesis | 3 | 48 | 0.0008 | 7.85E-03 | ko00905 |
| Diterpenoid biosynthesis | 4 | 209 | 0.0089 | 7.14E-02 | ko00904 |

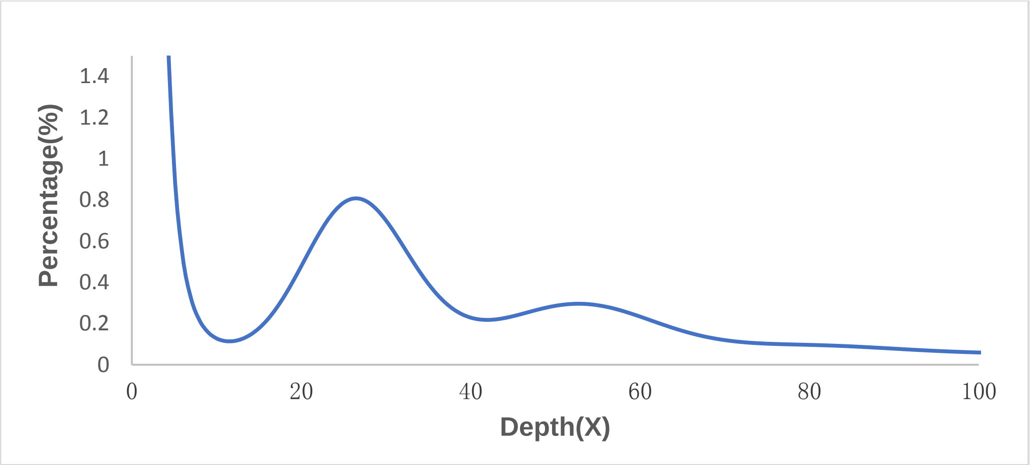


**Supplementary Fig. 1. The *k*-mer frequency distribution.** A total of 2.73 Tb clean sequencing data from 350 bp libraries were used to conduct *k*-mer analysis (*k* = 19). The genome size was estimated to be 82.55 Gb.

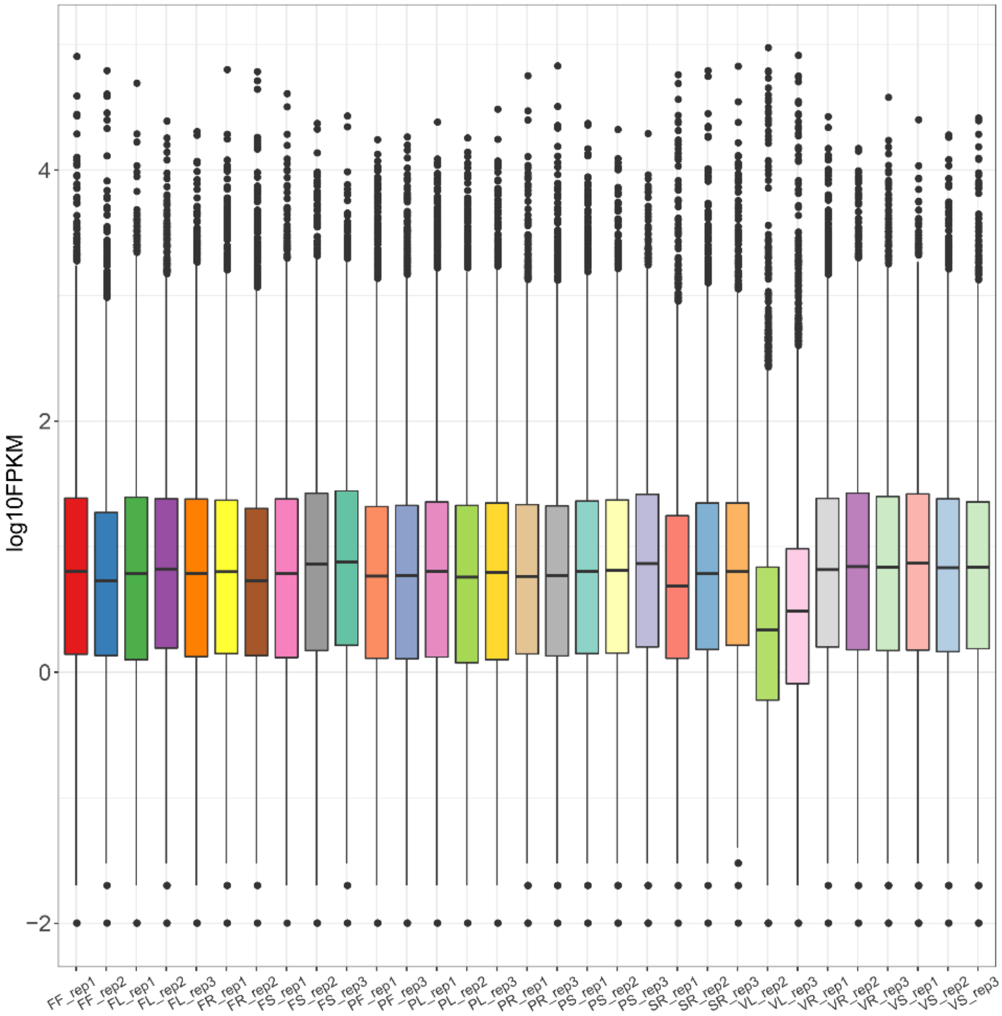


**Supplementary Fig. 2. Boxplot of gene expression.** X-axis refers to sample names and Y-axis refers to $\log_{10}$FPKM.

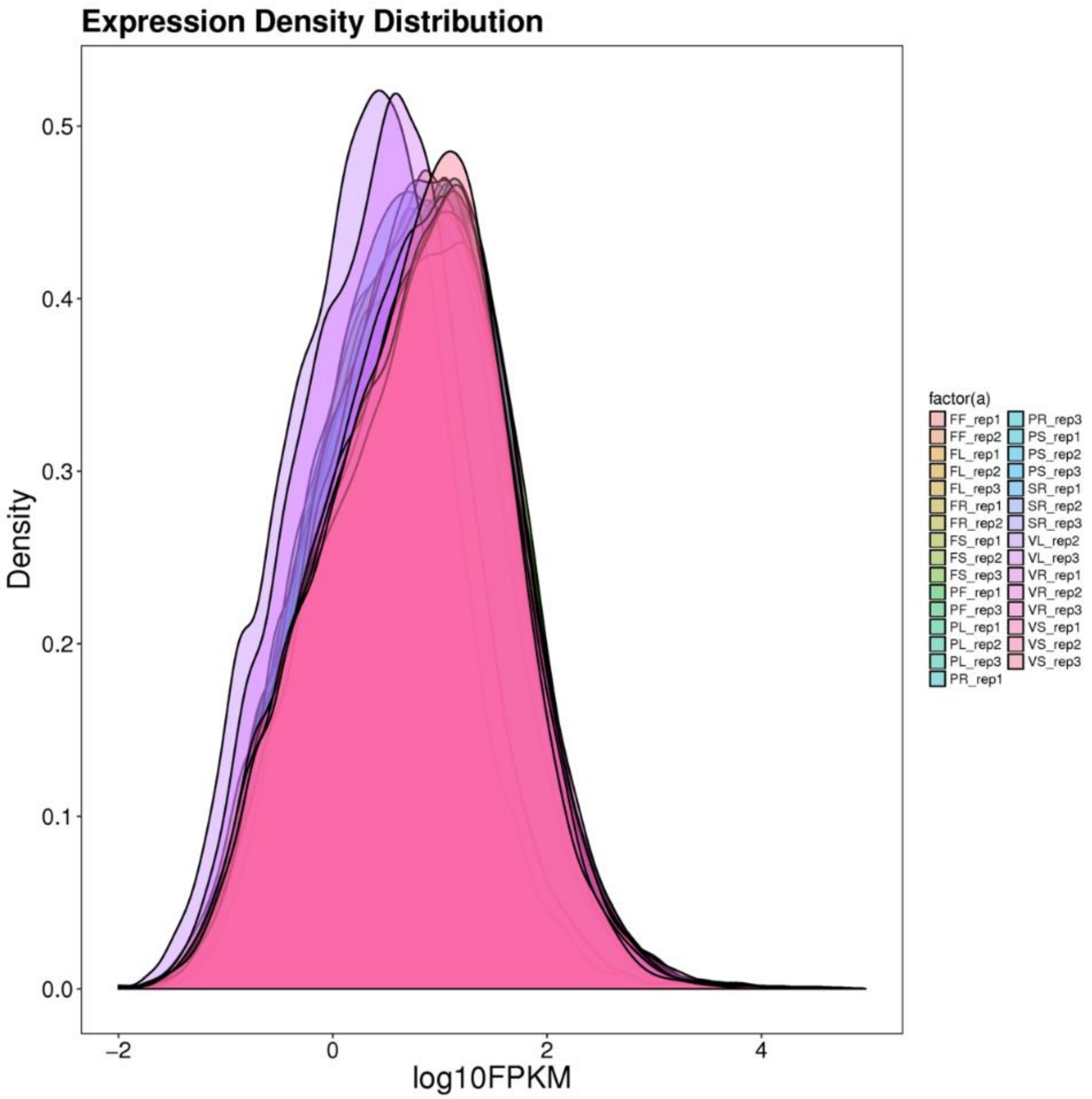


**Supplementary Fig. 3. Density distribution of gene expression.** X-axis refers to $\log_{10}$FPKM and Y-axis refers to the ratio of genes to the total number of expressed genes.

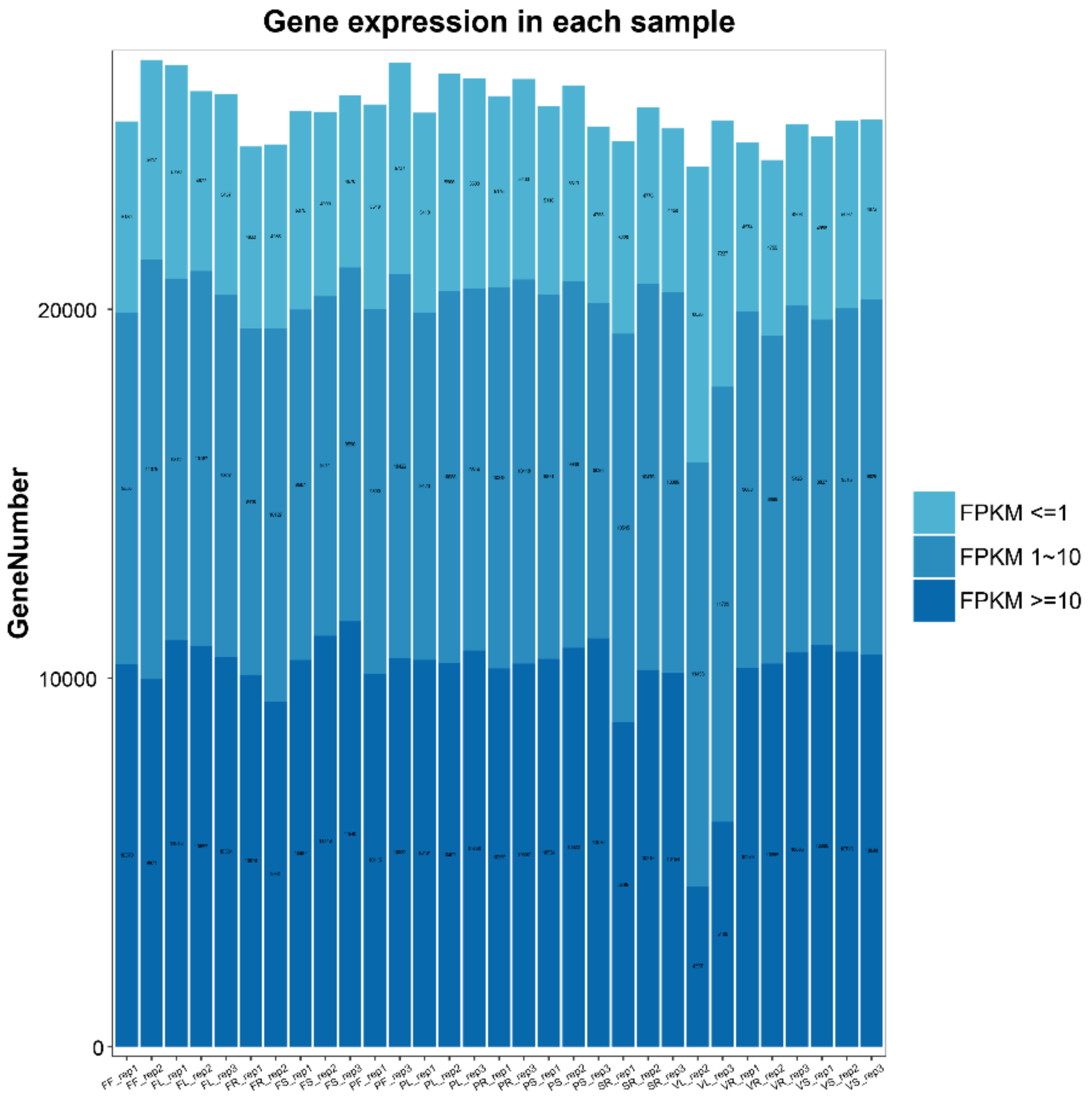


**Supplementary Fig. 4. Distribution of gene expression.** X-axis refers to sample names and Y-axis refers to gene numbers, the color shades indicate different levels of expression.

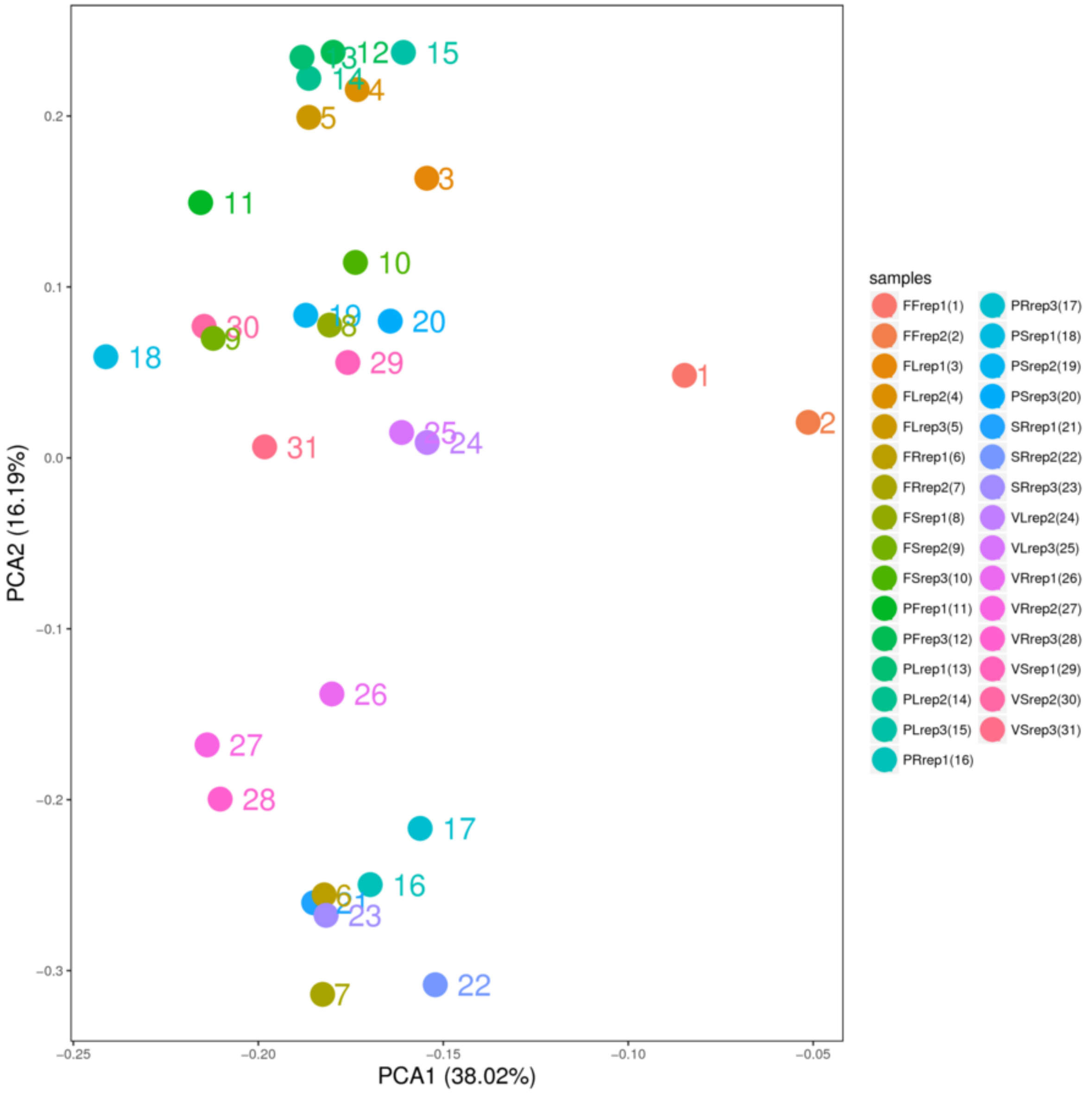


**Supplementary Fig. 5. PCA analysis of all the samples.** The X and Y axes represent the new data set of corresponding principal components obtained after the dimensionality reduction treatment of the sample expression, which is used to represent the distance between samples. The values in the coordinate axis label brackets represent the percentage of the variance of the corresponding principal component interpretation population.

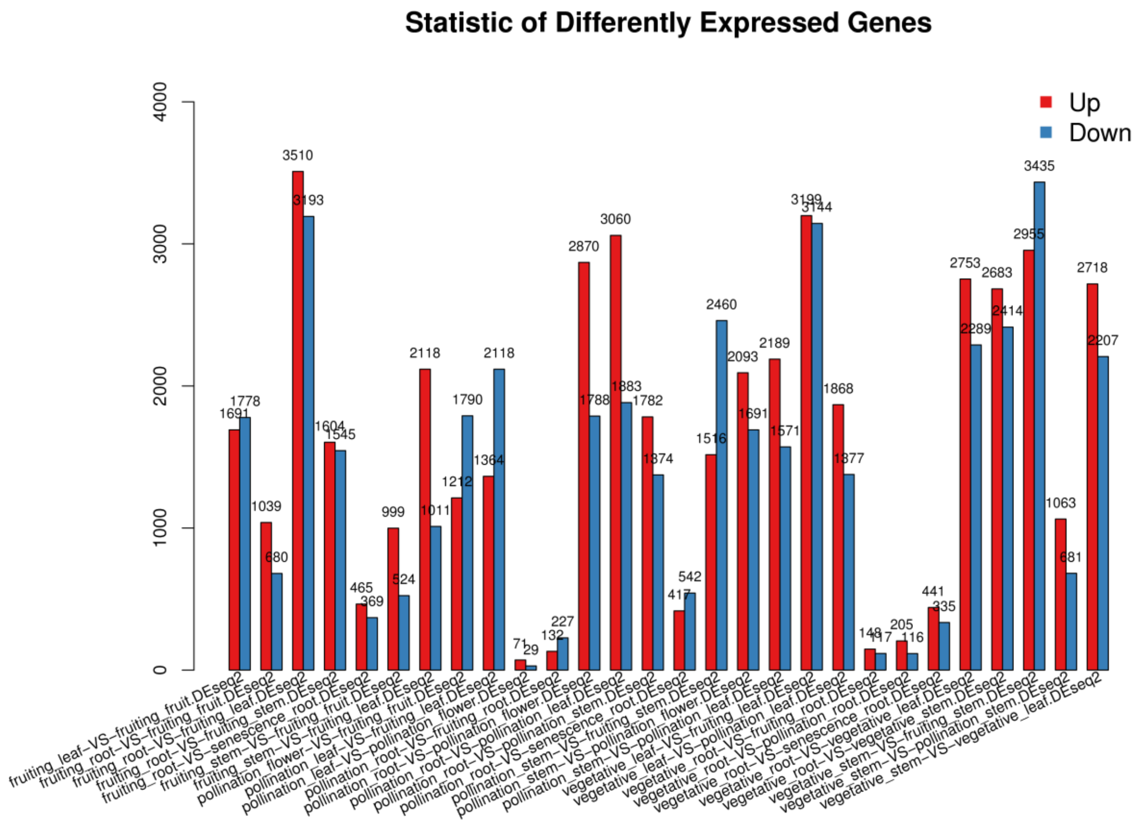


**Supplementary Fig. 6.** Statistics of DEG numbers between different tissues and growth stages.

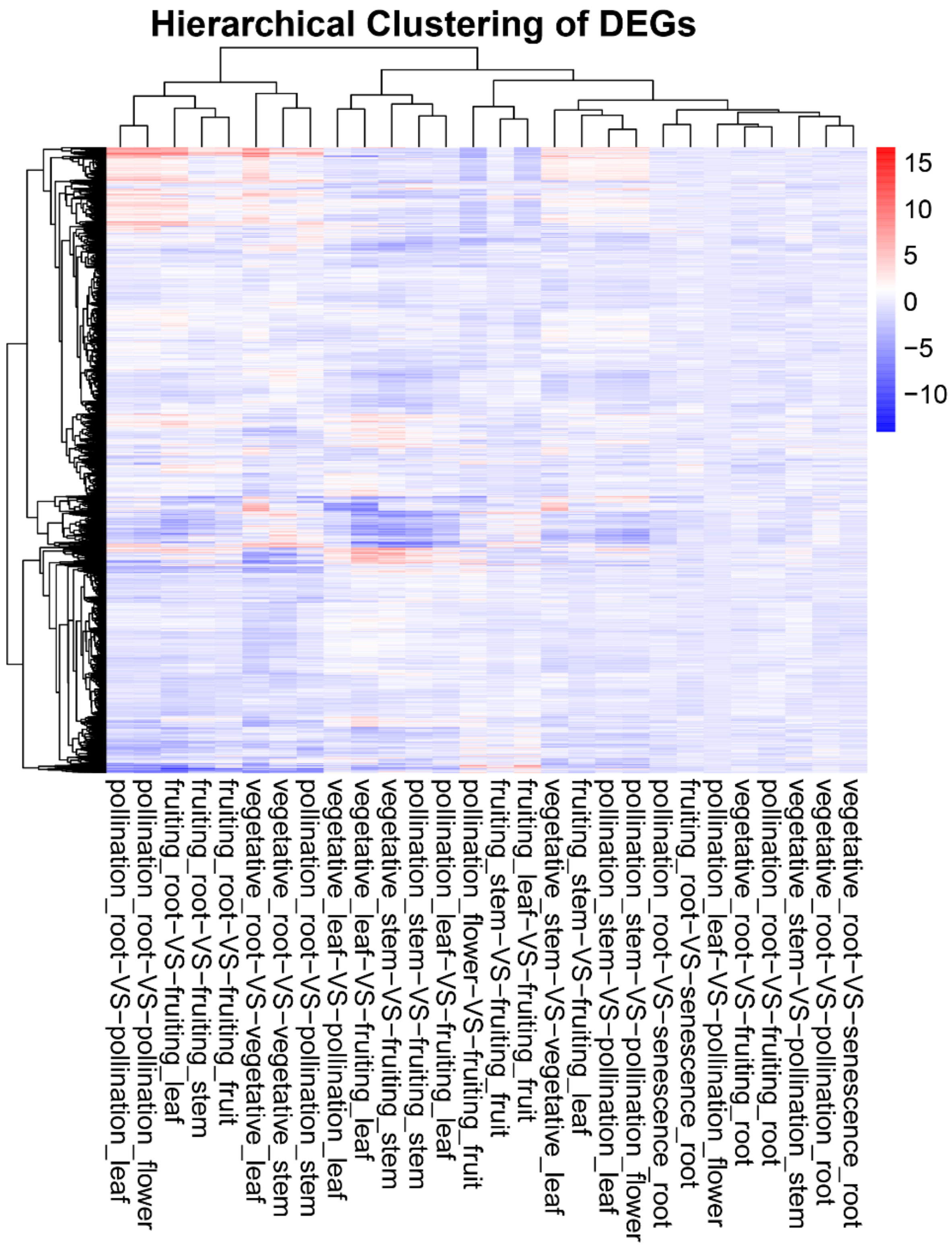


**Supplementary Fig. 7. Heatmap of DEG hierarchical clustering.** The X-axis represents the two groups undergoing cluster analysis, and Y-axis represents the significantly different genes.

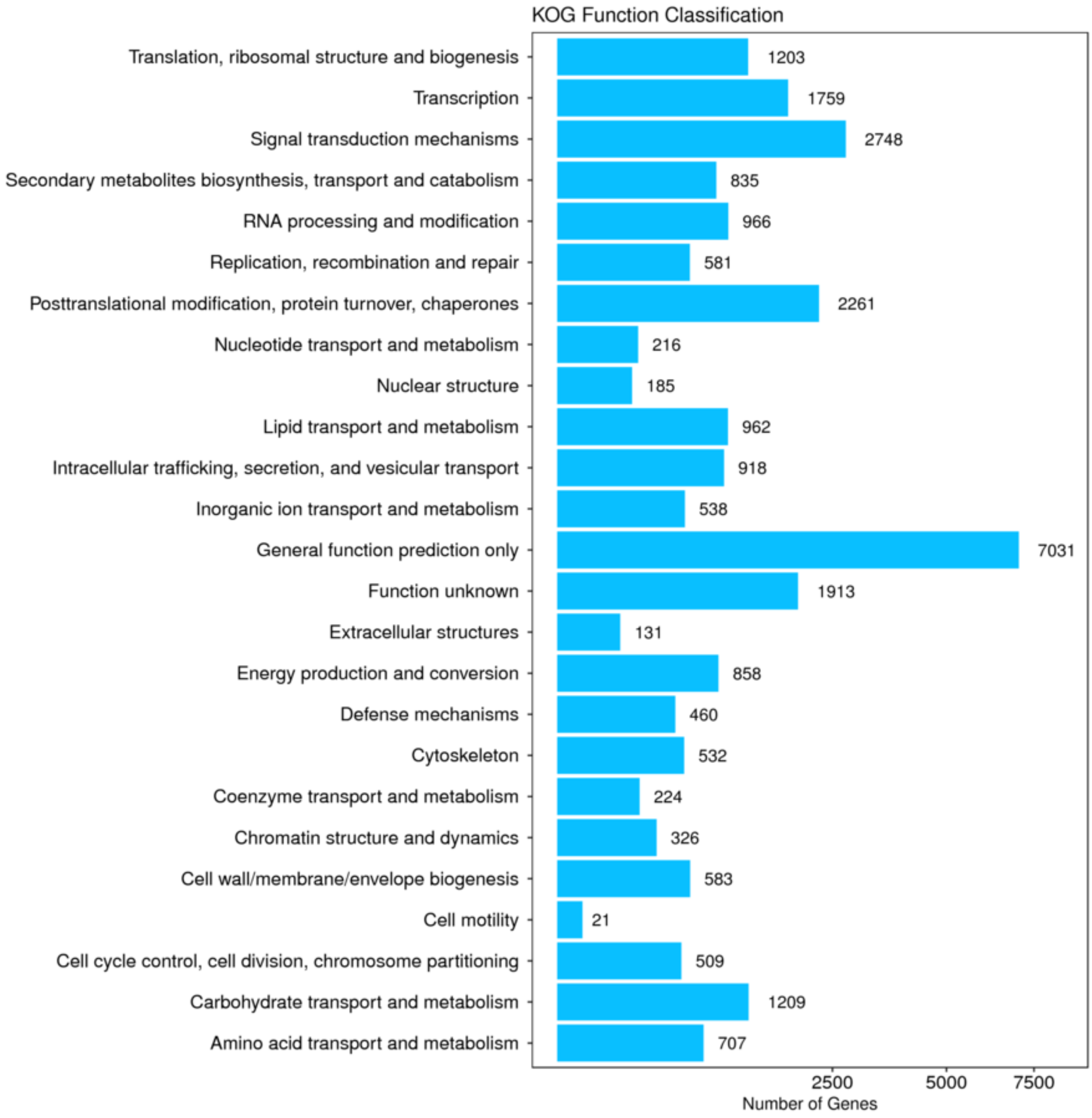


**Supplementary Fig. 8. Distribution of KOG function annotation.** X-axis refers to gene numbers and Y-axis refers to KOG function names.

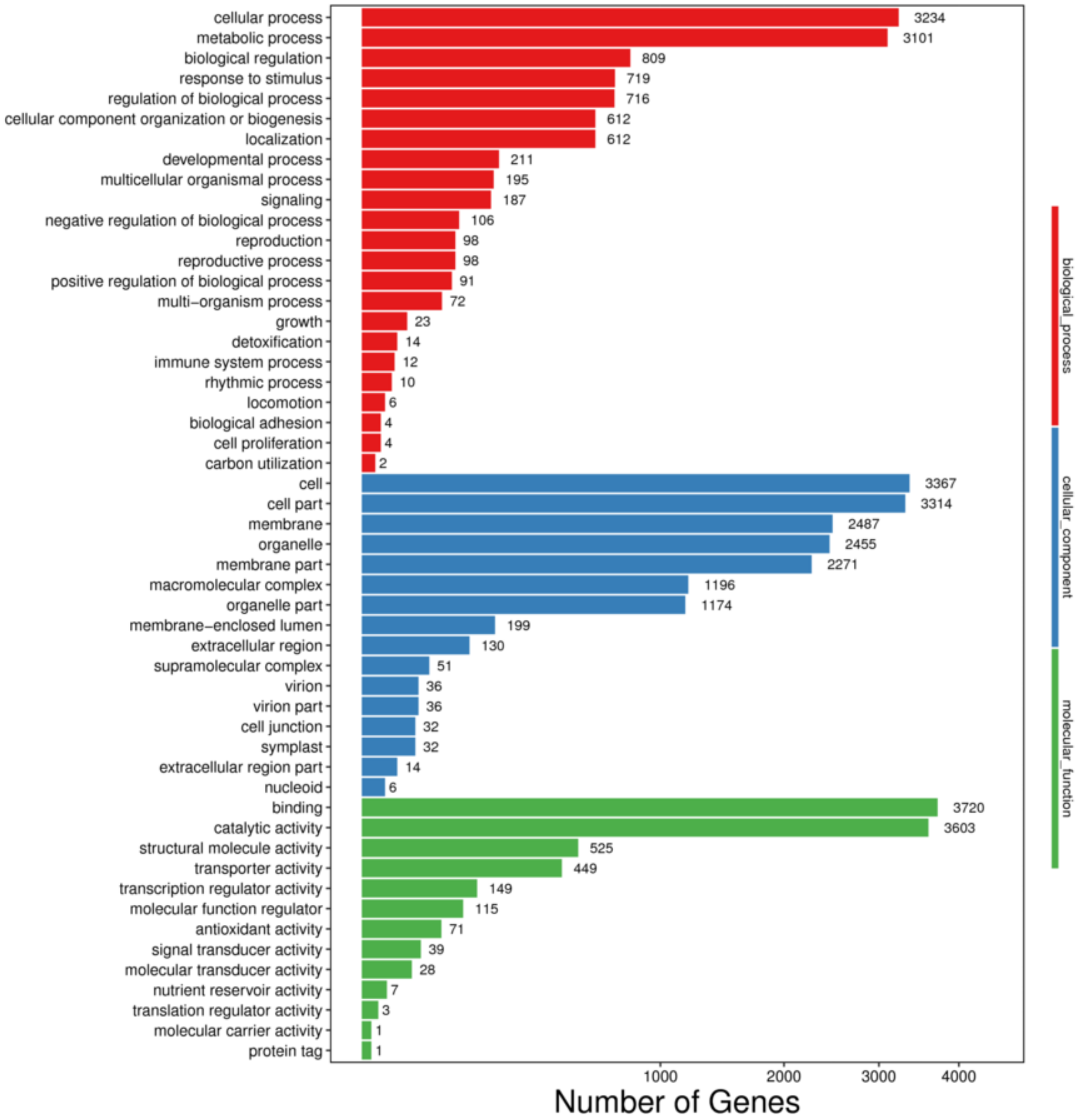


**Supplementary Fig. 9.** Distribution of GO function annotation. X-axis refers to gene numbers and Y-axis refers to GO function names.

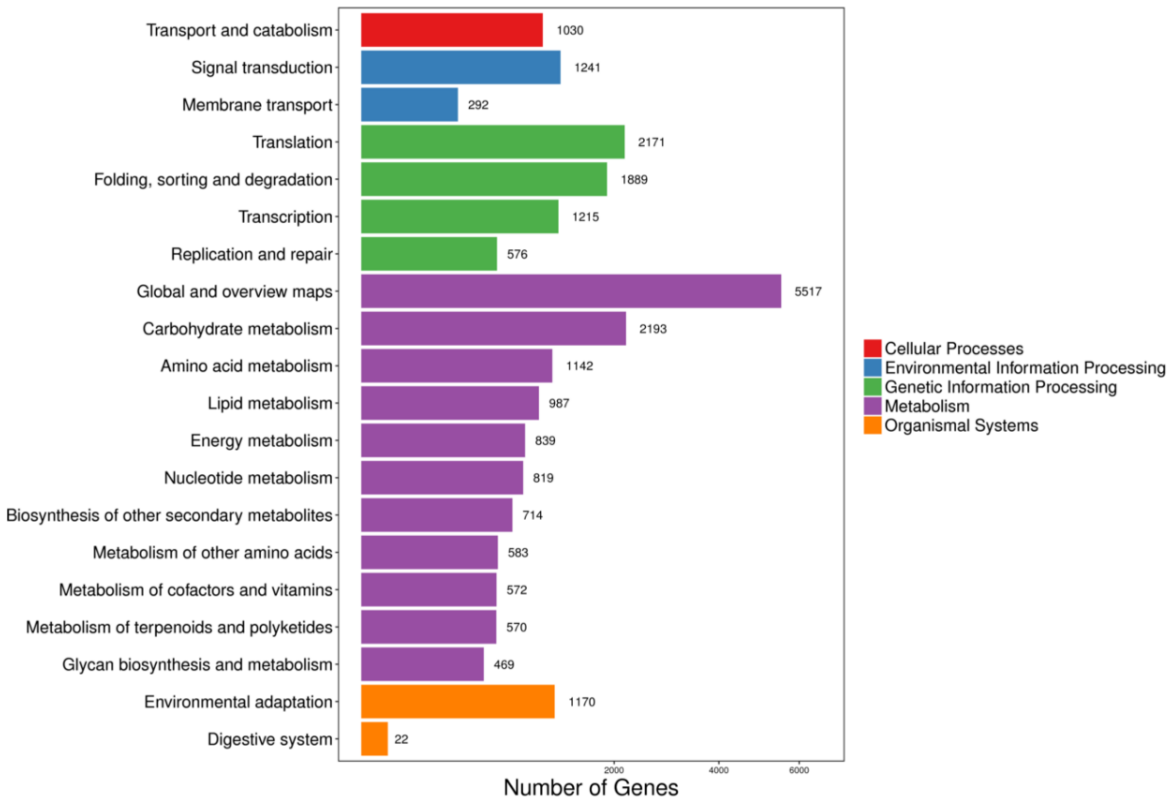


**Supplementary Fig. 10.** Distribution of KEGG function annotation. X-axis refers to gene numbers and Y-axis refers to KEGG function names.

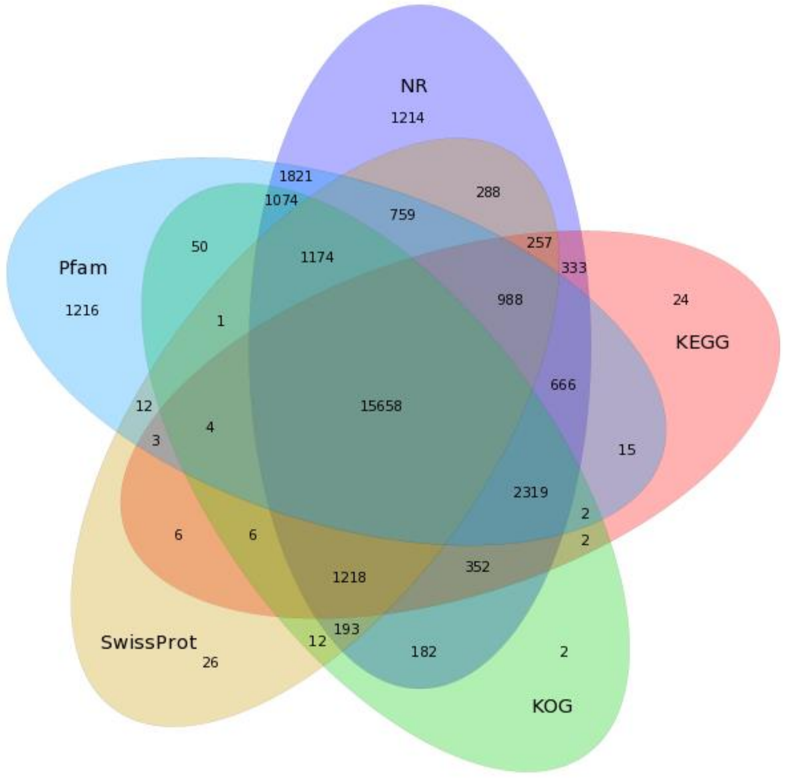


**Supplementary Fig. 11.** Venn diagram of function annotation in NR, KOG, KEGG, Pfam and SwissProt databases.

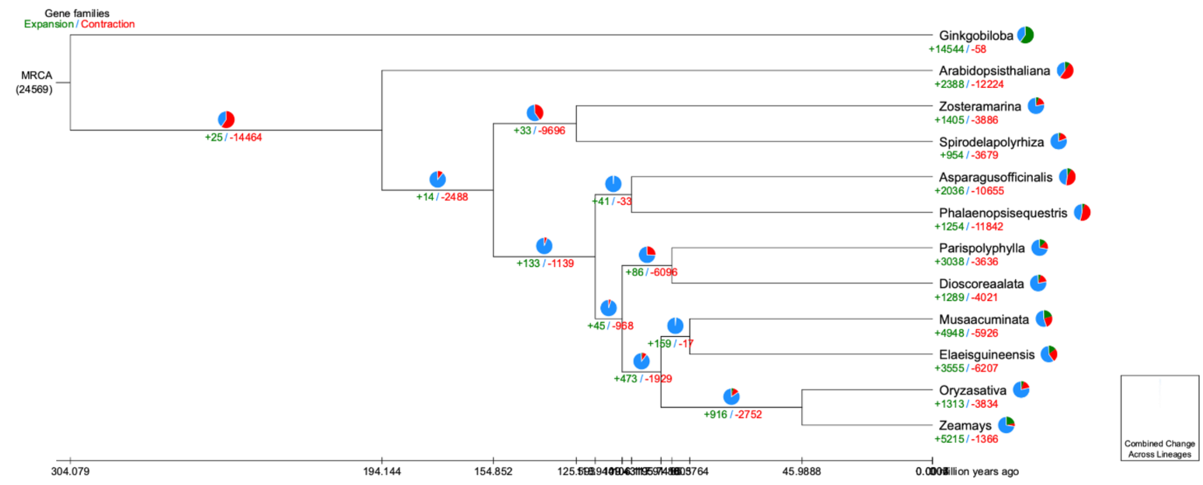


**Supplementary Fig. 12. The expanded and contracted gene families in PPY.**

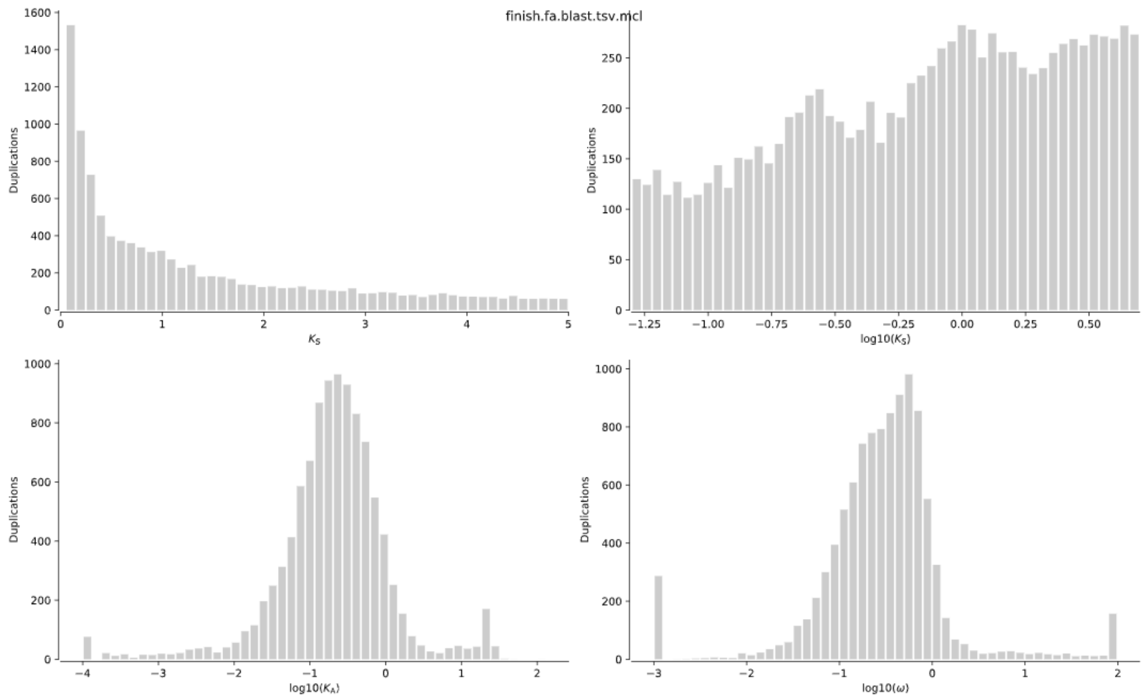


**Supplementary Fig. 13. The estimation of the whole genome duplication event of PPY.**

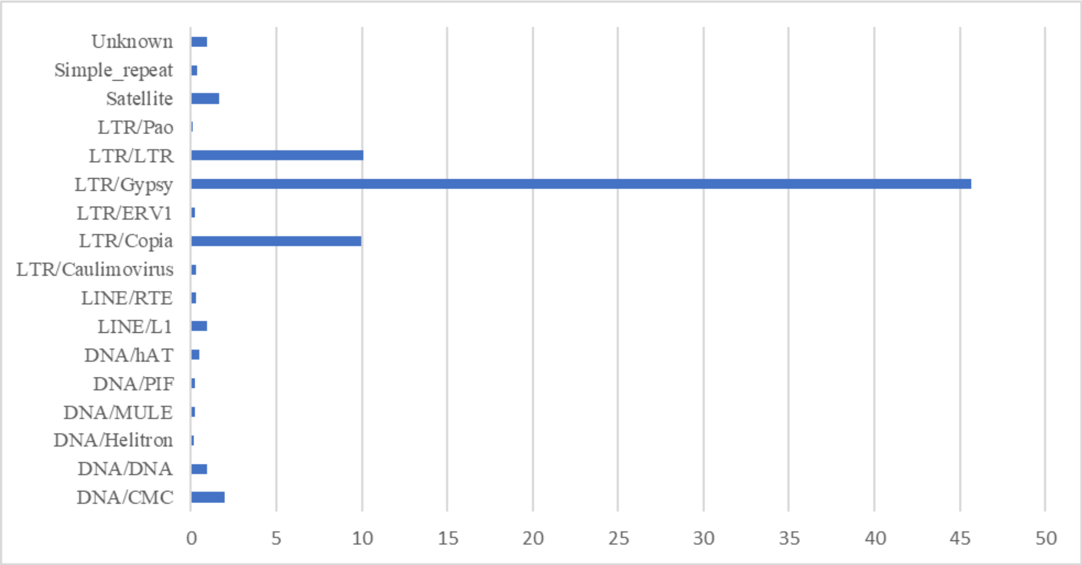


**Supplementary Fig. 14. Summary of the repetitive sub-type of PPY genome.**

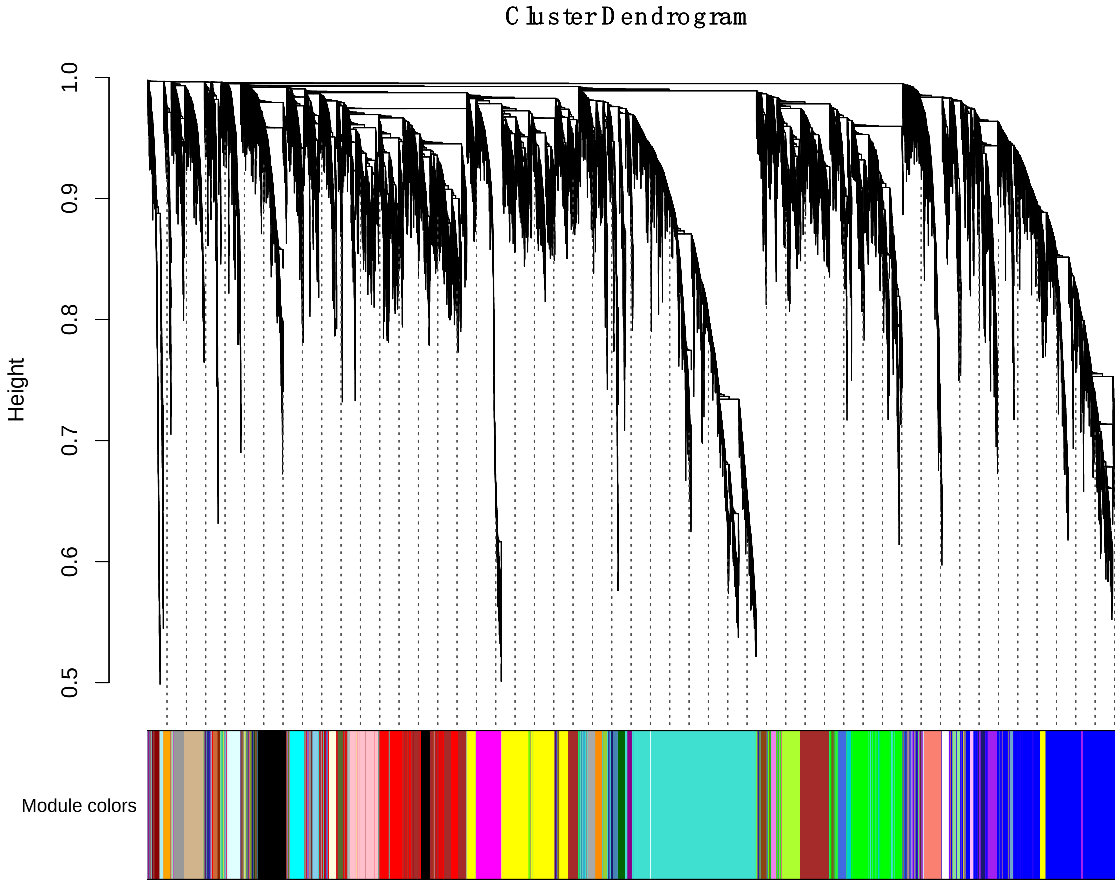


**Supplementary Fig. 15. The mode detection results of WGCNA analysis using 31 RNA-seq data.**

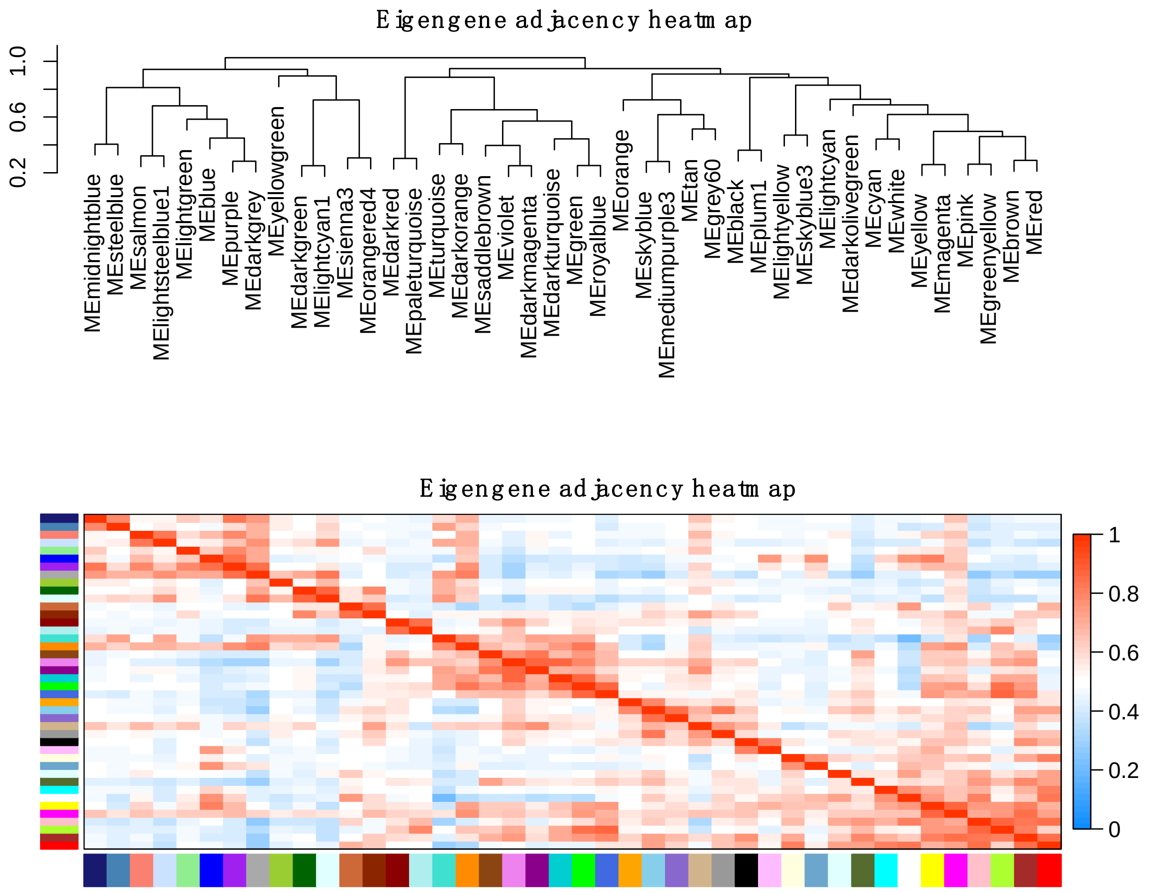


**Supplementary Fig. 16. The correlation analysis among different modes of WGCNA.**

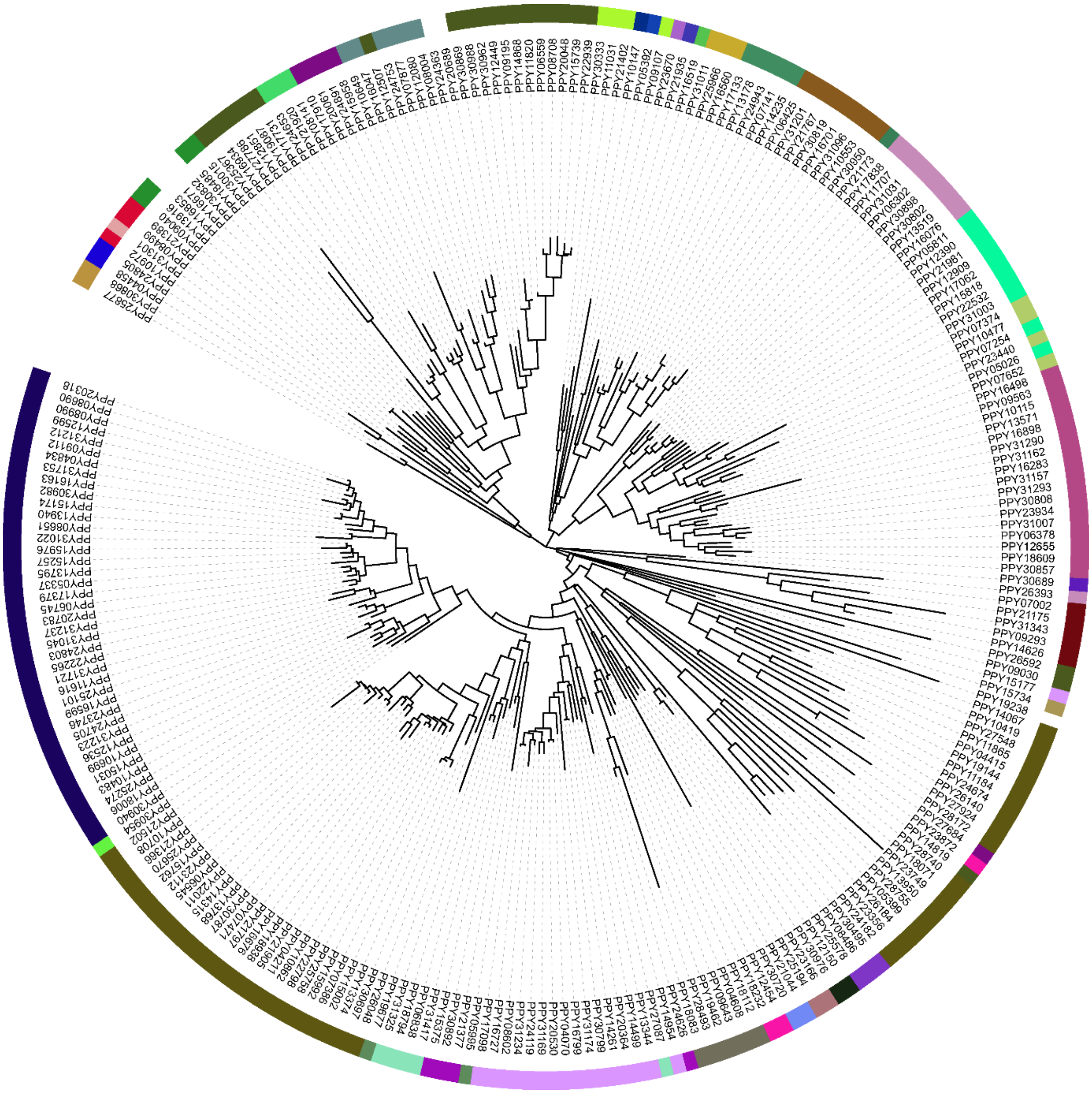

**Supplementary Fig. 17. The phylogenetic tree of the identified CYP genes of PPY.**

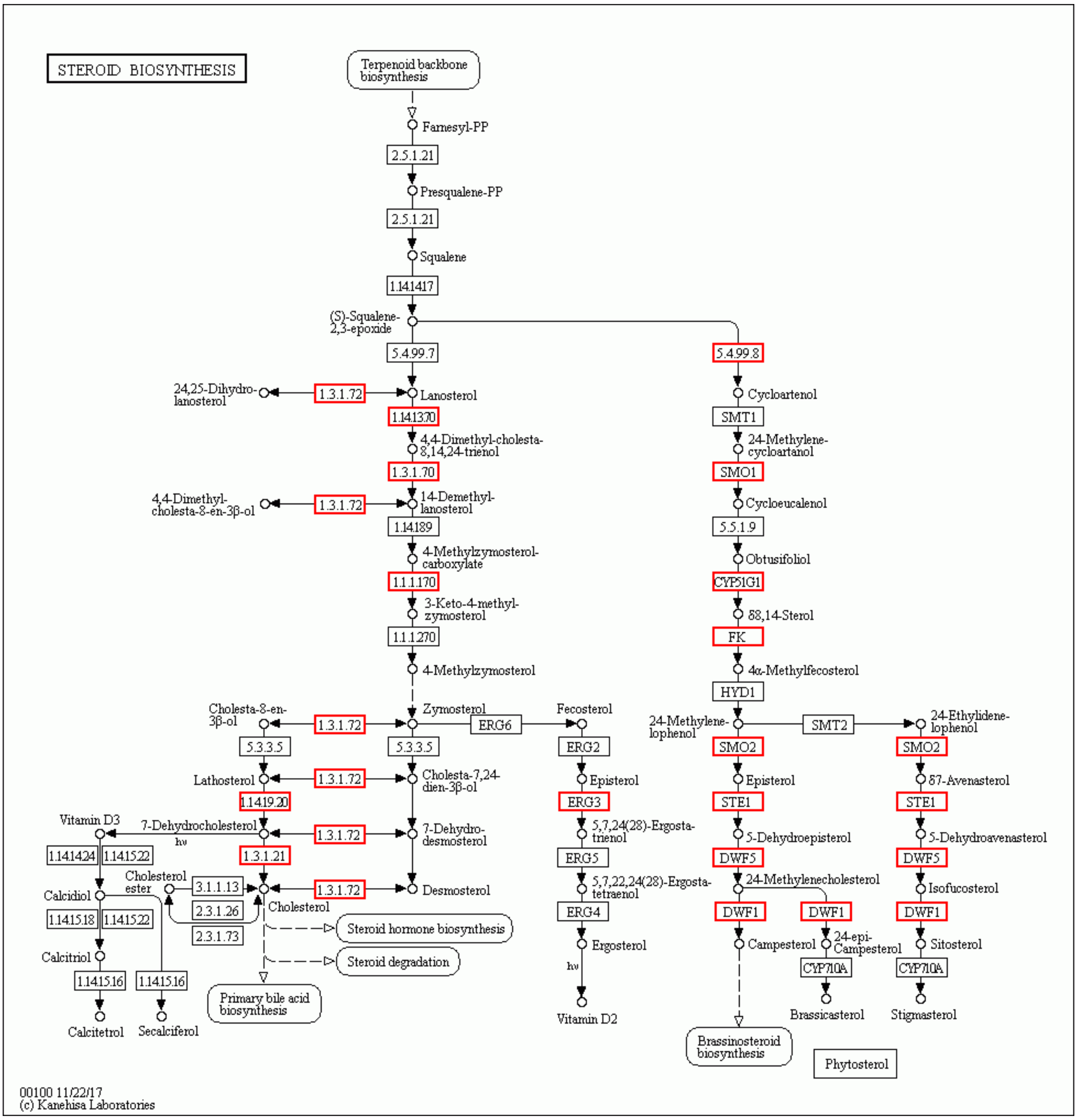


**Supplementary Fig. 18. Mapping of the genes in module 35 to KEGG pathways.** Homologues of the enzymes in red boxes can be found in PPY proteins.

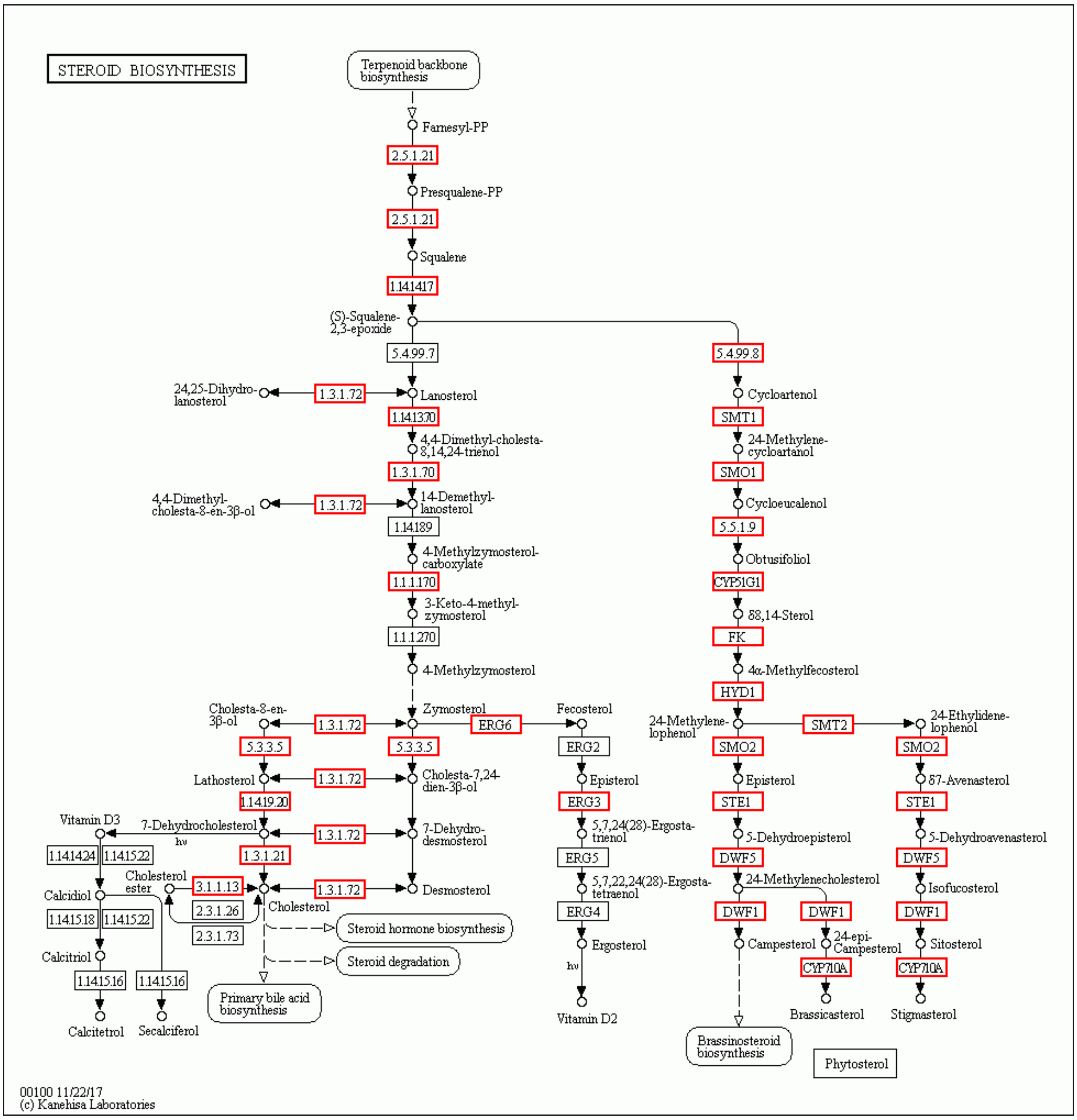


**Supplementary Fig. 19. Mapped KEGG steroid biosynthetic network.** Homologues of the enzymes in red boxes can be found in PPY proteins.
